# Dynamic PRC1-CBX8 stabilizes a porous structure of chromatin condensates

**DOI:** 10.1101/2023.05.08.539931

**Authors:** Michael Uckelmann, Vita Levina, Cyntia Taveneau, Xiao Han Ng, Varun Pandey, Jasmine Martinez, Shweta Mendiratta, Justin Houx, Marion Boudes, Hari Venugopal, Sylvain Trépout, Qi Zhang, Sarena Flanigan, Minrui Li, Emma Sierecki, Yann Gambin, Partha Pratim Das, Oliver Bell, Alex de Marco, Chen Davidovich

## Abstract

The compaction of chromatin is a prevalent paradigm in gene repression. Chromatin compaction is commonly thought to repress transcription by restricting chromatin accessibility. However, the spatial organisation and dynamics of chromatin compacted by gene-repressing factors are unknown. Using cryo-electron tomography, we solved the three-dimensional structure of chromatin condensed by the Polycomb Repressive Complex 1 (PRC1) in a complex with CBX8. PRC1-condensed chromatin is porous and stabilised through multivalent dynamic interactions of PRC1 with chromatin. Mechanistically, positively charged residues on the internally disordered regions (IDRs) of CBX8 mask negative charges on the DNA to stabilize the condensed state of chromatin. Within condensates, PRC1 remains dynamic while maintaining a static chromatin structure. In differentiated mouse embryonic stem cells, CBX8-bound chromatin remains accessible. These findings challenge the idea of rigidly compacted polycomb domains and instead provides a mechanistic framework for dynamic and accessible PRC1-chromatin condensates.

## Main

Chromatin structure is intricately linked to transcriptional activity^1^. Compacted or “closed” chromatin is generally associated with inhibition of transcription while “open”, more accessible chromatin is more prone to being transcribed^1^. Polycomb Repressive Complex 1 (PRC1) is a repressive chromatin modifier critical for organismal development^2,3^. PRC1 has been proposed to inhibit gene expression by tightly compacting chromatin in a process that often is considered to restrict chromatin accessibility^2–11^. However, direct evidence for PRC1-compacted chromatin being inaccessible is sparse and mechanistic explanations remain unsatisfactory (reviewed in^11^). Furthermore, recent studies show that changes in chromatin accessibility are more gradual than the simple binary classification into “open” and “closed” chromatin suggests^12–15^. A challenge in consolidating these seemingly contradictory findings is limited information into how PRC1 influences the three dimensional structure of chromatin.

PRC1 complexes can include one of five different chromobox proteins (CBX), all homologous to the fly Polycomb (Pc)^16^. The CBX protein CBX2 forms condensates through liquid-liquid phase separation, providing a potential mechanism for the compartmentalization of facultative heterochromatin^9,10^. Phase separation is emerging as a mechanism for chromatin organisation through the association of self-similar domains^17^. Chromatin can form condensates in the presence of divalent cations and histone tails^17–20^. Within these condensates, chromatin has variably been described as liquid-like, formed through liquid-liquid phase separation^17,19^, or as solid^20^. A recent structure of liquid-liquid phase-separated chromatin, condensed by magnesium cations without protein binding partners, revealed that nucleosomes organise into irregular assemblies^19^. The lack of apparent periodicity in chromatin geometry has also been noted in computational simulations and in first attempts to image chromatin in cells by cryo-electron tomography (cryo-ET)^21,22^. However, the structural arrangement of chromatin condensed by a repressive factor remained unknown.

Herein we describe the three dimensional cryo-ET structure of chromatin condensed by a polycomb-repressive complex. We focus on a PRC1 complex that includes CBX8 (PRC1^C8^), a chromobox protein that is upregulated during cell differentiation^23^ and has oncogenic potential^24,25^. We show that dynamic interactions between PRC1^C8^ and chromatin promote condensates through phase separation. Mechanistically, positive charges on the internally disordered regions (IDRs) of CBX8 are required for DNA binding and chromatin condensation. Contrary to expectations, PRC1-condensed chromatin is not tightly compacted but stabilises a porous chromatin structure that allows largely unhindered diffusion of PRC1^C8^.

## Results

### PRC1-chromatin condensates are porous and accessible

To determine the structure of polycomb-compacted chromatin and the mechanisms of polycomb-driven chromatin compaction, we reconstituted the system *in vitro*. The reconstitution included a chromatinized polycomb target gene (3,631 bp DNA) with a sequence from the human ATOH1 locus, which can harbour roughly up to 20 nucleosomes, assuming 150-200 bp per nucleosome. This construct is referred to as chromatin hereafter. We used a native DNA sequence for chromatin reconstitution, as regular spacing using artificial nucleosome stabilising sequences were previously reported to drive the liquid-liquid phase separation of chromatin^17^. The nucleosomes on the chromatin that we reconstituted are not evenly phased (Extended Data Fig. 1a). The purified recombinant PRC1 complex is composed of RING1B, BMI1 and CBX8 (PRC1^C8^) (Fig. 1a). The PRC1^C8^ complex is pure (Fig. 1b), monodispersed (Fig. 1c) and retains H2A ubiquitylation activity comparable to the RING1b-BMI1 heterodimer (Fig. 1d). This also applies to all other protein complexes used in this study (Extended Data Fig. 2 and 1b,c).

**Fig. 1.**
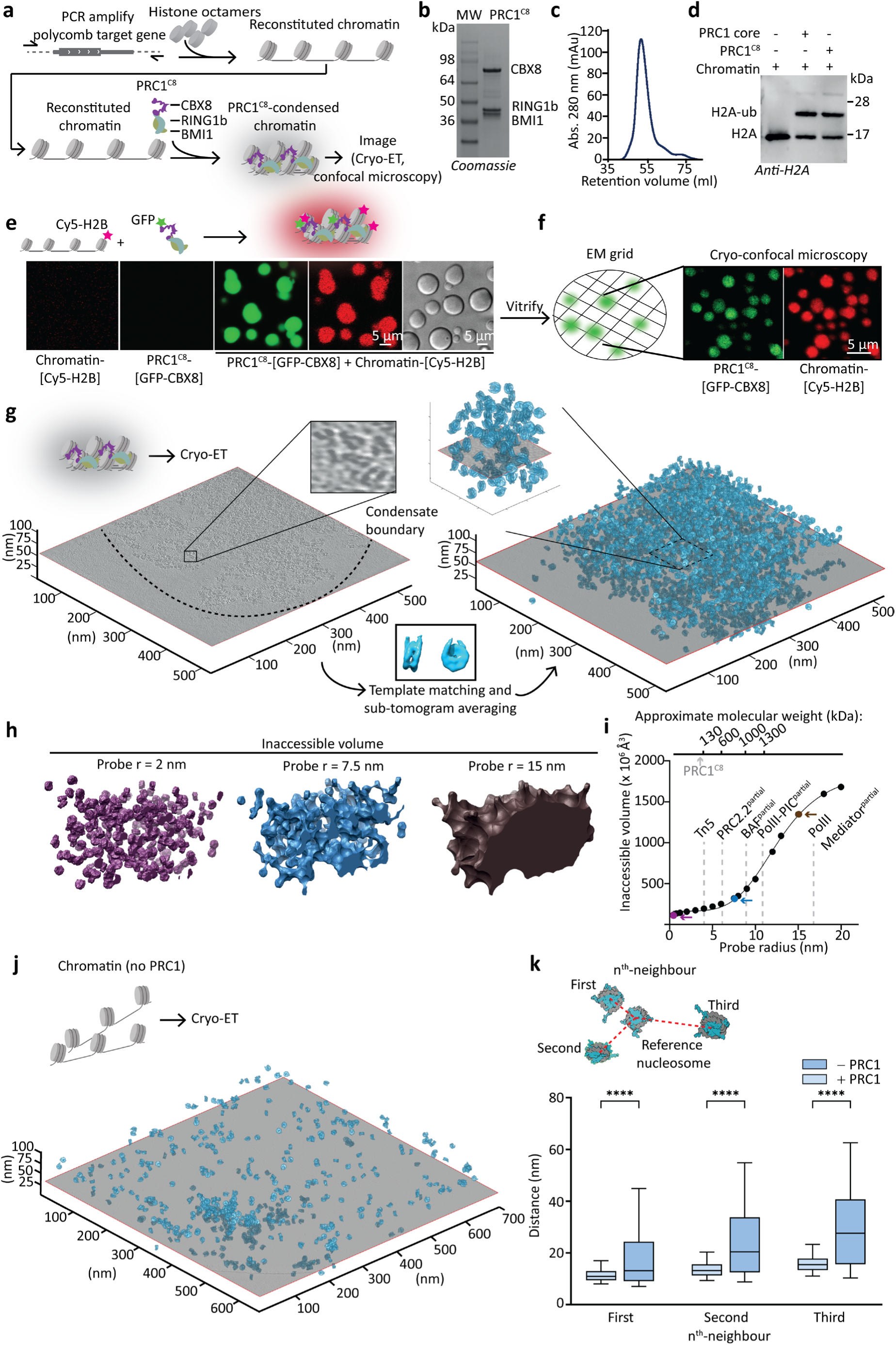
The molecular structure of PRC1-chromatin condensates is porous and accessible to macromolecules. **a,** Schematics introducing the workflow. **b,** SDS-PAGE of purified PRC1^C8^ complex that includes RING1b, BMI1 and MBP-tagged CBX8. **c,** Size exclusion chromatography of the purified PRC1^C8^ using a HiLoad Sephacryl 300 16/60 column. **d,** In vitro ubiquitylation assay comparing PRC1^C8^ to a RING1b-BMI1 heterodimer. Samples in lane 2 and 3 included E1, E2, ubiquitin and ATP. Ubiquitylation is detected by western blot using an anti-H2A antibody. **e,** Chromatin condensates induced by the PRC1^C8^ complex and the individual proteins, visualised by confocal (left and centre) and phase contrast (right, from independent experiments) microscopy. CBX8 is GFP-labelled and chromatin is Cy5 labelled. Protein and chromatin concentrations are 1 μM and 20 nM (estimated nucleosome concentration), respectively. **f,** Cryo-confocal microscopy of vitrified PRC1^C8^-chromatin condensates. **g,** Cryo-electron tomography of a PRC1^C8^-chromatin condensate. Shown is a central slice through the reconstruction (left image). Nucleosome subtomogram averages (centre, bottom) are then placed in a volume the size of the tomographic slice, at the position and orientation determined by template matching and subtomogram averaging (right image). 1330 nM PRC1^C8^ and 3500 nM chromatin (estimated nucleosome concentration) were assayed in 3.5 mM HEPES-KOH pH 7.5, 6.8 mM TRIS-HCl PH 7.5, 21 mM NaCl, 7 mM KCl, 0.8 mM DTT. See Supplementary Data 1 for a list of cross correlation peaks. **h,** Surface representation of the volume of a subset of the PRC1^C8^-chromatin condensate structure that is inaccessible to probes of given radii. **i,** Inaccessible volumes for a given probe radii are plotted, with exemplary molecules indicated (in grey) and selected probes coloured as in **h**. For the indicated complexes, the hydrodynamic radius was estimated^32^ using resolved domains from published structures (see Methods section for PDB accessions) as a minimum size estimate. **j,** As in g but without PRC1^C8^. See Supplementary Data 2 for a list of cross correlation peaks. **k**, Pairwise distances of each individual nucleosome to its nearest three neighbouring nucleosomes in 3D space in tomograms with and without PRC1^C8^. Only unique pairs are plotted from two tomograms with (+PRC1) and three tomograms without (-PRC1). Whiskers extend from the 5th to the 95th percentiles. Significance was tested using a Brown-Forsythe ANOVA with a Games-Howell post hoc test. **** = p-value < 0.0001.

When combined, chromatin and PRC1^C8^ were sufficient to form spherical phase-separated condensates, apparent in differential interference contrast (DIC) and fluorescence imaging (Fig. 1e). Using two different fluorescence labels, we confirmed the presence of both chromatin and PRC1^C8^ within the same condensates (Fig. 1e). Importantly, both PRC1^C8^ and chromatin are necessary for chromatin condensation, while the individual components do not phase-separate (Fig. 1e). PRC1^C8^-chromatin condensates are preserved on an EM grid after vitrification (Fig. 1f). To study these structures by cryo-electron tomography, we reduced the salt concentration and increased chromatin concentration (see Methods section). This was done to decrease condensate size and abundance, which was necessary for high-quality data collection (Fig. 1g). We used a PRC1 complex with MBP-tagged CBX8 for cryo-tomography. The MBP tag does not affect condensation, as condensates still form after tag cleavage (Extended Data Fig. 1d). We collected tomograms near the borders of condensates, to observe the boundary conditions (Extended Data Fig. 3a for an example of a condensate on the grid). Subtomogram averaging allowed us to identify the orientation and position of individual nucleosomes in the tomographic volume (Extended Data Fig. 3b-d).

The reconstruction reveals a dense network of hundreds of nucleosomes with a distinct condensate boundary (Fig. 1g and Movie S1). We could not unambiguously assign density to PRC1^C8^, possibly because it adapts various conformations while simultaneously using multiple surfaces to interact with chromatin (more below). The final structure reflects the arrangement of nucleosomes in PRC1^C8^-chromatin condensate (Fig. 1g, second panel). Unexpectedly, the structure shows that PRC1^C8^ does not compact nucleosomes into an impassable barrier. Instead, PRC1^C8^ rather stabilises chromatin in a porous mesh-like structure (Fig. 1g). Analysing the orientation of individual nucleosomes towards their neighbouring nucleosomes shows no obvious orientation bias (Extended Data Fig. 3c,d). We conclude that PRC1^C8^ does not induce a substantial inter-nucleosome orientation bias, but rather supports forming a porous chromatin structure.

We next wished to determine the size of macromolecules that could diffuse into PRC1-chromatin condensates. We used the condensate structure to calculate solvent-excluded volumes^26^ with variable probe radii ranging from 0.2 nm to 20 nm (Fig. 1h,i). Interestingly, the analysis shows that the condensate is accessible for macromolecules of a considerable size of up to 8 nm in radius (equivalent to approximately 600 kDa). Small macromolecules (<10 kDa), with radii below 2 nm, would have enough room to access every single nucleosome. Conversely, access is increasingly restricted for molecules with a radius above 8 nm (approximately 600 kDa). This suggests that PRC1-chromatin condensates are surprisingly accessible and that PRC1^C8^ itself would be able to move within these condensates largely unhindered.

To compare the structure of PRC1^C8^-chromatin condensates to PRC1^C8^-free chromatin, we generated cryo-tomograms of chromatin without PRC1^C8^ (Fig. 1j, movie S2). Low magnification cryo-EM images show that condensation happens only in the presence of PRC1^C8^ (Extended Data Fig. 4a,b) and condensates are preserved on the EM grid. The structure of chromatin in the absence of PRC1^C8^ is less dense than in the presence of PRC1^C8^, with distances between neighbouring nucleosomes that are on average significantly longer (Fig. 1j,k). These results confirm that PRC1^C8^ facilitates large-scale chromatin restructuring.

We observe some sporadic areas of high nucleosome density, even without PRC1 (Extended Data Fig. 4c,d). The median distance to the next neighbouring nucleosomes in these sporadic dense PRC1^C8^-free condensates are very similar to distances measured in the PRC1^C8^-chromatin condensates (9.8 nm and 10.9 nm, respectively, Extended Data Fig. 4d). This matches closely to distances reported for chromatin condensed by MgCl2, where the radial distribution function of nucleosomes peaked at 10.6 nm^19^. At the nuclear periphery in cells, the median distance between neighbouring nucleosomes is about 12 nm, which is again remarkably similar^27^. Overall, this raises the possibility that PRC1 thermodynamically stabilises a naturally-occurring condensed chromatin state, rather than actively compacts chromatin. By doing so, PRC1 may cause multiple compacted arrays to cluster together more often.

### PRC1^C8^ is mobile and chromatin is static within PRC1-chromatin condensates

To gain further insights into PRC1^C8^-chromatin condensation, we tested the dependency of condensation on the concentration of PRC1 and chromatin. Condensates form under close to physiological monovalent salt concentrations of 122.5 mM (90 mM KCl and 32.5 mM NaCl), at PRC1^C8^ concentrations as low as 250 nM and are dependent on PRC1^C8^ (Fig. 2a). At a high concentration of PRC1^C8^ (2,000 nM), most efficient condensation occurs at high chromatin concentration (850 nM nucleosome concentration). Yet, at lower concentration of PRC1^C8^ (250 nM), ideal condensation appears at lower chromatin concentration and the condensation efficiency is then reduces when the chromatin concentration increases (Fig 2a). This is possibly because at high chromatin concentration the large amount of potential binding sites for PRC1^C8^ reduces the average per-site-occupancy of PRC1^C8^. This may lead to less efficient phase separation. A similar observation was recently made for a PRC1 complex with CBX2 and PHC2^28^. Altering the salt concentrations confirms that PRC1^C8^-chromatin condensates most readily form close to physiological salt concentration (Extended Data Fig. 5). We conclude that PRC1^C8^ is sufficient to drive the formation of the chromatin condensates under physiologically relevant conditions. Furthermore, the efficiency of phase separation depends on the ratio of PRC1 to chromatin. We next asked if chromatin and PRC1 show different dynamics within the condensates.

**Fig. 2.**
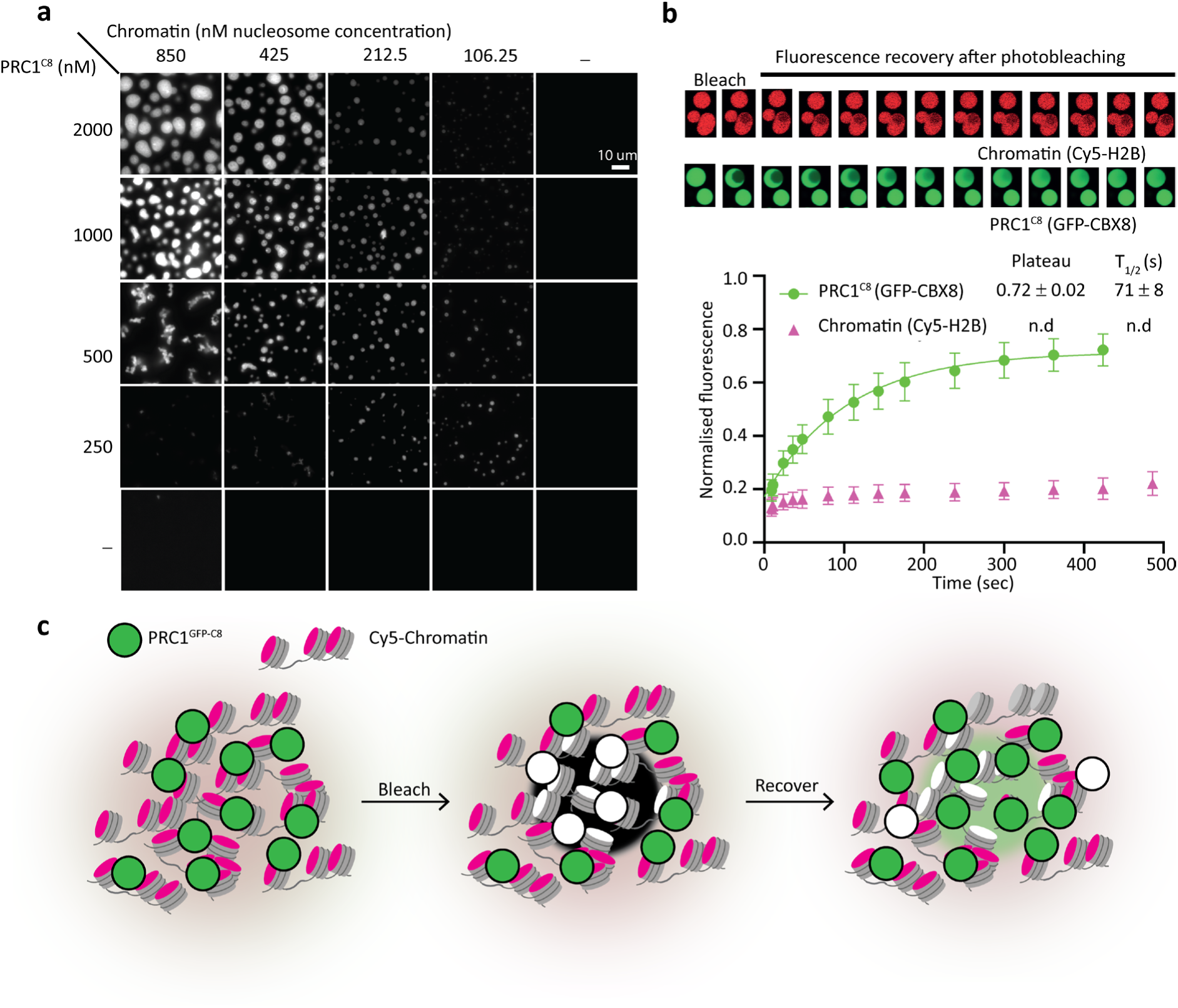
PRC1^C8^ is mobile while chromatin is static within PRC1-chromatin condensates. **a,** Titration of Chromatin against PRC1^C8^. Condensates were assessed with a fluorescence widefield microscope imaging a Cy5 label on the chromatin. Presented are representative micrographs of three replicates, including two with MBP-tagged PRC1^C8^ (presented) and one with GFP-tagged PRC1^C8^. **b,** Representative micrographs of FRAP recorded in PRC1^C8^-chromatin condensates. CBX8 is GFP labelled and chromatin is Cy5 labelled. Mean fluorescence intensity of the bleached area, normalised to pre-bleach mean signal, is plotted for every time point. Error bars show standard deviation from n=7 (GFP) and n=8 (Cy5) measurements recorded from two independent experiments. The GFP signal recovery was fit with an exponential association model, best fit values for Plateau and fluorescence recovery half time (T_1/2_) are shown with standard error. **c,** Schematic representation of the FRAP experiment.

Fluorescence recovery after photobleaching (FRAP) of PRC1^C8^-chromatin condensates shows fast recovery kinetics for GFP-labelled PRC1 (Fig. 2b, in green; T_1/2_ = 71 ± 8 s). Conversely, we observed a very slow recovery for Cy5-labelled chromatin (Fig. 2b, in red). The GFP signal does not recover up to 100 % but rather plateaus at 72 ± 2%. This is possibly because a substantial percentage of the condensate has been bleached, while redistribution of GFP-labelled PRC1^C8^ within the same condensate likely drives fluorescence recovery during the monitored timescale. However, we cannot exclude the possibility that incomplete recovery is due to an immobile fraction of CBX8. To exclude the possibility that a large protein tag on CBX8 prevents it from remaining static on chromatin we used a synthetic fluorescence dye to sparsely label random lysine residues of PRC1^C8^. We then compared FRAP recovery for two different PRC1 complexes: one complex included CBX8 with an N-terminal MBP tag and the other included an untagged CBX8 (Extended Data Fig. 6a,b). The PRC1^C8^ with untagged CBX8 is still dynamic (Extended Data Fig. 6a, T_1/2_ = ∼1300 s), albeit recovery is slower when compared to MBP-tagged PRC1^C8^ (Extended Data Fig. 6b, T_1/2_ = 362 ± 29 s) and the GFP-tagged protein (Fig. 2b; T_1/2_ = 71 ± 8 s). Chromatin remains static in all samples. These results indicate that although the tagging and labelling strategies affect the dynamics of the system quantitatively, its overall behaviour remains qualitatively the same: within condensates, PRC1 is mobile while chromatin itself is static (Fig. 2c and Extended Data Fig. 6). This confirms that PRC1^C8^ can diffuse within PRC1^C8^-chromatin condensates, in line with our structural analysis (Fig. 1h,i). Recent publications have been at odds as to the solid or liquid state of chromatin^17,20^. Our data suggests that chromatin behaves as a solid-like material when condensed by PRC1^C8^.

We next wished to gain insights into PRC1^C8^ -chromatin condensate formations at low protein concentrations that better resemble physiological concentrations. To this end, we employed a single-molecule confocal microscope that allows tracking of individual condensates through a confocal volume^29^ (Extended Data Fig. 7a,b). We used GFP-labelled PRC1^C8^, where GFP peaks indicate the formation of bright protein assemblies. Importantly, this system allows the detection of protein assemblies smaller than what can be identified by standard fluorescence microscopes^30^. In the presence of chromatin, assemblies are observed at PRC1^C8^ concentrations as low as 62.5 nM (Extended Data Fig. 7c). This indicates that condensates can form at physiologically relevant PRC1 concentrations, previously estimated as 130 nM in polycomb bodies in cells^31^. In the absence of chromatin, GFP peaks are only detected sporadically, even at the highest PRC1^C8^ concentration (Extended Data Fig. 7 b,c). Overall, this data indicates that PRC1^C8^-chromatin condensates form under physiologically relevant PRC1^C8^ concentration, but do not form without chromatin.

### Multivalent interactions between PRC1^C8^ and chromatin induce phase separation

A scaffold-client based phase separation model has recently been proposed for PRC1-CBX2 complexes^28,35^, where CBX2^35^ or chromatin^28^ act as a scaffold that induces phase separation of PRC1 proteins. Since PRC1^C8^ is insufficient to phase-separate without chromatin (Fig 1e), we wished to test if chromatin might act as a scaffold that concentrates PRC1^C8^ and induces phase separation. To test this model, we probed the different interaction sites between PRC1 and chromatin. The whole PRC1^C8^ complex (RING1b, BMI1 and CBX8) is necessary and sufficient to condense chromatin, while the PRC1 core or CBX8 alone do not condense chromatin (Fig. 3a). This suggests multivalent interactions between the PRC1^C8^ complex and chromatin, involving different chromatin interacting surfaces in both PRC1 and CBX8. To identify the different interaction sites, we first used crosslinking mass spectrometry (XL-MS) to probe for protein-protein interactions within PRC1^C8^ –chromatin condensates (Fig. 3b and Supplementary Data 4; PRC1 with an MBP-tagged CBX8 was used). As expected, we identified extensive crosslinks between the RING domains of RING1B and BMI1. RING1B and BMI1 did not crosslink to histones. This is likely because these proteins bind to the acidic patch of the nucleosome^36^, which is unlikely to be crosslinked by the BS3 crosslinker that reacts preferentially with lysine residues. The results show multiple crosslinks from the CBX8 chromodomain to the H3 histone tail (Fig. 3b), indicative of binding. Interactions between CBX-proteins and H3K27me3-modified H3-histone tails have been proposed to recruit PRC1 to chromatin modified by PRC2^2,3^. However, CBX8 did crosslink to unmethylated H3 tails (Fig. 3b) and a trimethyl-lysine analogue (MLA) at H3K27 did not improve the chromatin-condensation activity of PRC1^C8^ (Fig. 3d). We conclude that H3K27me3 is not necessary for the chromatin condensation activity of PRC1^C8^ and that the H3 histone tail, even if unmodified, provides an interaction site for PRC1 on chromatin.

**Fig. 3.**
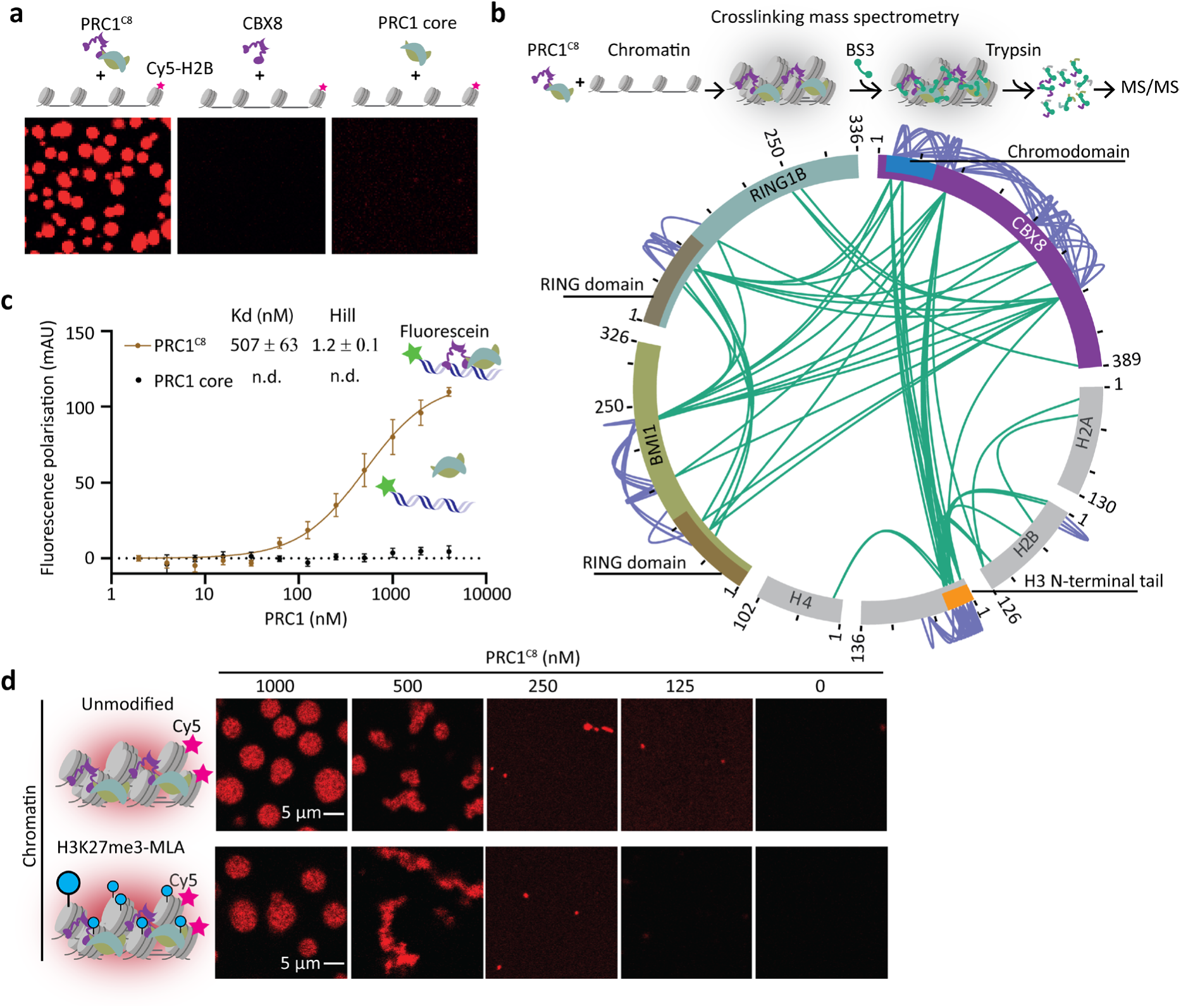
Multivalent interactions between PRC1^C8^ and chromatin. **a,** Chromatin condensation in response to the whole PRC1^C8^ complex (RING1b, BMI1 and CBX8) or the individual components CBX8 and the RING1b–BMI1 heterodimer. Representative images from two replicates. **b,** Intramolecular (purple lines) and intermolecular (green lines) protein-protein interactions mapped within PRC1-chromatin condensates using crosslinking mass spectrometry (XL-MS). Data is from three independent replicates. **c,** PRC1^C8^ or PRC1 core binding to a fluorescein labelled 24bp DNA probe measured by fluorescence polarisation. Data points show the mean (baseline subtracted) of three independent replicates and the error bars indicate the standard error. The continuous line represents the fit to a Hill binding model, when applicable. **d,** Titration of PRC1^C8^ to unmodified chromatin (top) and H3K27me3-MLA chromatin (bottom) at an identical chromatin concentration (50 ng/μl DNA) and 150 mM KCl. Micrographs are representative of two independent replicates.

We also observe extensive self-crosslinks in the IDRs of CBX8 (Fig. 3b). This may indicate inter- or intramolecular interaction within this flexible lysine-rich region. However, these self-crosslinks within the IDRs appear even in the absence of chromatin, suggesting that they are not related to condensation (Extended Data Fig. 8 and Supplementary Data 3). DNA has previously been shown to bind CBX8^37^ and could provide another interaction site for PRC1^C8^ on chromatin. We tested DNA binding in solution using a 24bp double stranded DNA probe and found that CBX8 is necessary for the DNA-binding activity of PRC1^C8^ (Fig. 3c). Hence, CBX8 binding to DNA provides a second interaction site of PRC1 with chromatin.

We conclude that PRC1 interacts with chromatin via at least three distinct sites: PRC1^C8^ binds to DNA and the H3 tail via CBX8, as shown herein (Fig. 3), and binds the nucleosome acidic patch via RING1b-BMI1 as shown elsewhere^36^. These multivalent interactions would have to change dynamically while PRC1 maintains the condensed state of chromatin and diffuses through it at the same time (Fig. 2b). Collectively, we propose that PRC1 induces chromatin condensation via phase separation, through dynamic multivalent interactions between PRC1 and chromatin.

### DNA binding by the CBX8 IDRs is required for efficient phase separation

To test how different PRC1^C8^-chromatin interaction sites affect phase separation, we generated several different chromatin and CBX8 mutants (Fig. 4a and Extended Data Fig. 9). Removing the CBX8 chromodomain (PRC1^C8ΔChromo^), which interacts with the H3 tail^37,38^ (Fig. 3b), does not have a significant effect on phase separation (Fig. 4b,c). We then mutated 21 positively charged residues in the CBX8 IDRs to alanine (PRC1^C8KR21A^). PRC1^C8KR21A^ is analogous to a phase separation-deficient CBX2 mutant that was previously studied^10^. Accordingly, PRC1^C8KR21A^ shows a clear defect in phase separation activity (Fig. 4b,c).

**Fig. 4.**
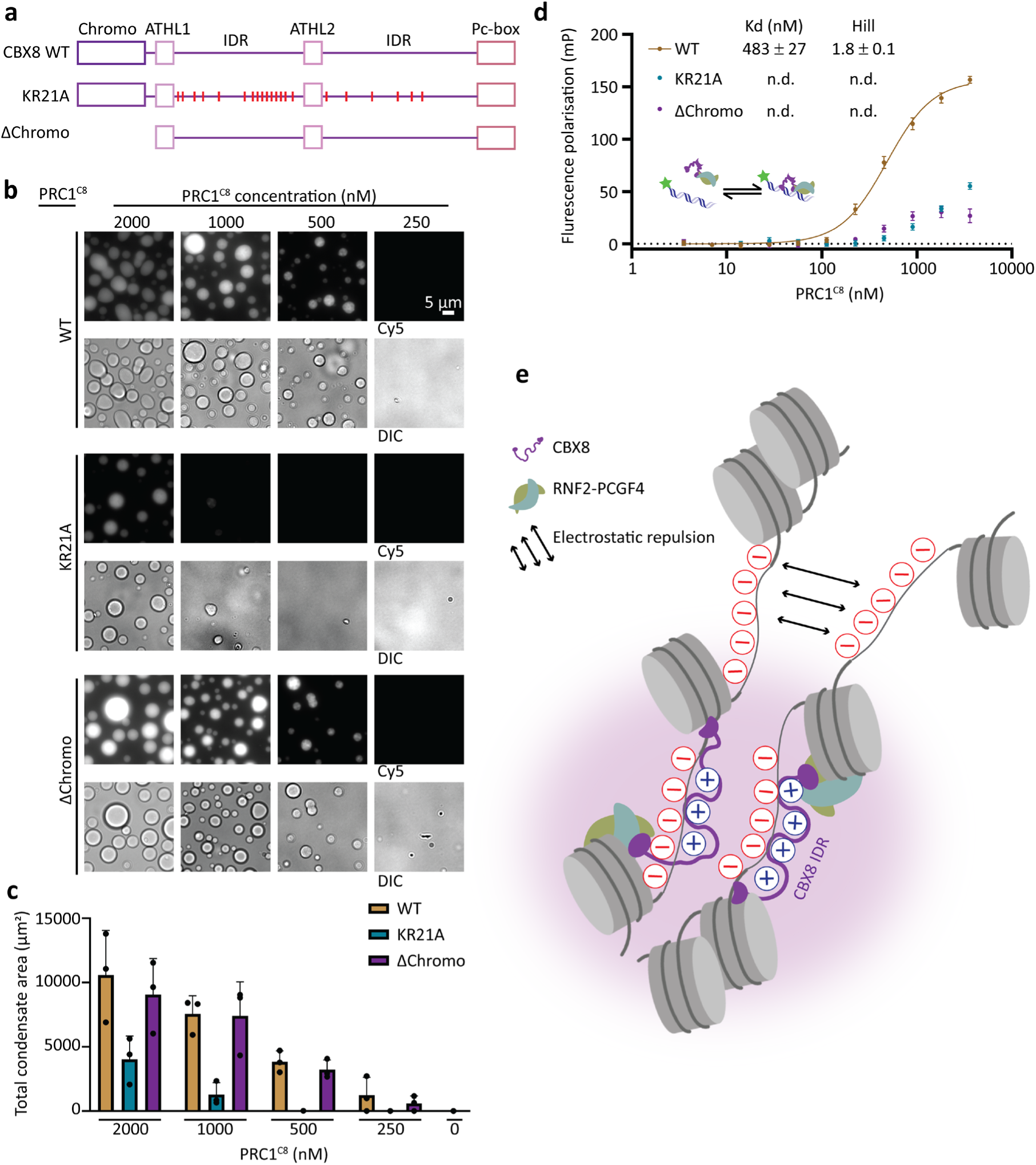
Positive charges in the CBX8 IDRs are required for DNA binding and phase separation. **a,** Schematics depicting the different CBX8 mutants used, drawn to scale. **b,** PRC1^C8^-chromatin condensates in the context of different CBX8 mutants. Varying concentrations of PRC1^C8^ were titrated to a constant concentration of C5-labelled reconstituted chromatin (50 ng/μl). Widefield fluorescence and differential interference contrast (DIC) micrographs are representative of three replicates. **c** Quantification of the total area covered by condensates per micrograph for different PRC1^C8^ mutants and concentrations. Bars represent the means from three independent replicates and error bars represent the standard deviation. **d,** Fluorescence polarisation assay measured the affinity of different PRC1^C8^ mutants for a Fluorescein-labelled 24 bp DNA probe. Data points are the mean (baseline subtracted) of three independent replicates and error bars indicate the standard error. The continuous line represents the fit to a Hill binding model, when applicable. **e,** Model for chromatin condensation by PRC1^C8^: Electrostatic interaction between the CBX8 IDR with DNA provides charge screening and promotes phase separation.

To further dissect the mechanism, we tested the DNA binding activity of PRC1^C8KR21A^ and PRC1^C8ΔChromo^ against a 24 bp double-strand DNA probe with a sequence from a polycomb-target gene. Both mutants are defective in DNA binding (Fig 4d). Hence, while DNA binding of the chromodomain was reported previously^37^, our data now suggests that the IDRs of CBX8 are also contributing to DNA binding. We hypothesise that electrostatic interactions between the negatively charged DNA to positive charges in the IDR lead to charge screening, which promotes phase separation of chromatin (Fig. 4e).

We next set to determine if the H2A acidic patch of the nucleosome affects PRC1^C8^-chromatin condensation. For that, we mutated residues in the H2A acidic patch that were previously shown to interfere with the interaction between PRC1 and the nucleosome^36^ (Extended Data Fig. 9). We observed a change in the condensate morphology (Extended Data Fig. 9a): while PRC1^C8^-chromatin condensates appear spherical, the condensates with mutated acidic patch chromatin adapt elongated and branched structures (Extended Data Fig. 9a; most apparent at lower PRC1^C8^ concentrations). The effects of the PRC1^C8KR21A^ mutant and the acidic patch chromatin mutant are additive (Extended Data Fig. 9b), suggesting that both interaction sites contribute to phase separation independently. The PRC1^C8ΔChromo^ mutant does not substantially affect phase separation, regardless of the chromatin used (Extended Data Fig. 9c). Overall, our data supports a model where PRC1^C8^ uses its chromodomain and IDRs to bind DNA. Then, PRC1^C8^ condenses chromatin through interactions between the IDRs and DNA and, independently, between PRC1 and the acidic patch on the nucleosomes (Fig 4e).

### CBX8 binding sites on chromatin in mouse embryonic stem cells are accessible

Given the porous structure of PRC1^C8^ -condensed chromatin in vitro (Fig. 1) and the dynamic diffusion of PRC1^C8^ within condensates (Fig. 2b), we next wished to probe for the accessibility of PRC1^C8^ -bound chromatin in cells. We carried out the Assay for Transposase Accessible Chromatin (ATAC-seq) in differentiated mESC, combined with ChIP-seq for CBX8 and H3K27me3. We used differentiated mESC, because CBX8 is expressed at very low levels in pluripotent mESC and is upregulated during retinoic acid-induced cell differentiation (^23^ and Fig. 5a). The DNA-loaded Tn5 used in ATAC-seq experiments forms a dimeric complex of approximately 130 kDa with an estimated hydrodynamic radius^39^ of 4.6 nm (based on PDB code 1MUH^40^). In agreement with the accessibility analysis *in vitro* (Fig. 1h), the majority of CBX8 ChIP-seq peaks in cells overlapped with ATAC-seq peaks (Figure 5b), indicating they are accessible to Tn5. This observation was persistent across the genome, where ATAC-seq peaks are co-localised with CBX8 and H3K27me3 peaks (Fig. 5c), indicating that CBX8-target genes are largely accessible. The insufficiency of CBX8 to restrict chromatin accessibility is further supported by the similar ATAC-seq profiles of wildtype and *Cbx8* knockout mESCs (Fig. 5c,f, compare blue to orange). Hence, although the overall chromatin accessibility is reduced during mESC differentiation (Fig. 5d,e), in agreement with previous works,^41^ this process is not dependent on CBX8. Collectively, we show that CBX8-bound polycomb-repressed chromatin is largely accessible in differentiated mESCs (Fig. 5b-f).

**Fig. 5.**
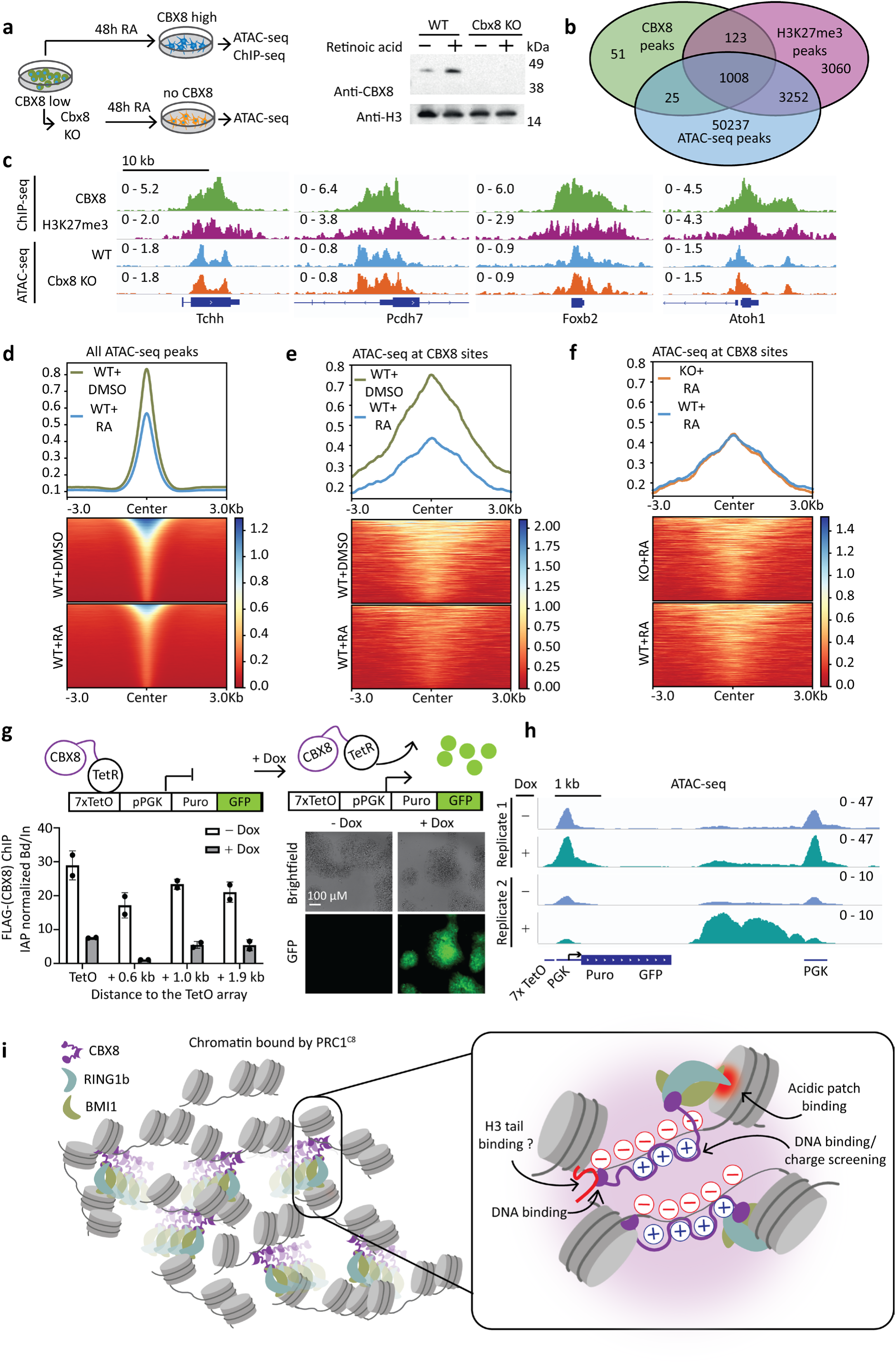
CBX8 binding sites on chromatin in mouse embryonic stem cells are accessible. **a,** Schematics of the experimental setup (left) and anti-CBX8 western blot (right) of wildtype and *Cbx8* knockout mESC after 48 hours of retinoic acid (RA) treatment. **b,** Overlap of ATAC-seq peaks and CBX8 and H3K27me3 ChIP-seq peaks. ATAC-seq peaks are defined from two biological replicates. **c,** ChIP-seq traces for H3K27me3 and CBX8 in wildtype mESC and representative ATAC-seq traces at four genes in wildtype and CBX8 knockout mESC after 48 hours of RA treatment. **d,** Accessibility changes at all ATAC-seq peaks in wildtype (WT) mESC, in response to retinoic acid (RA) treatment. **e,** Accessibility changes at all CBX8-target sites in wildtype mESC, in response to RA treatment. **f,** Comparison of accessibility at CBX8-target sites between wildtype and *Cbx8* knockout cells after RA treatment. **g,** Top: schematic representation of the chromosome-integrated reporter. Left panel: ChIP-qPCR using FLAG antibody (CBX8 is FLAG tagged) at indicated distances from the TetO array, in the presence and absence of doxycycline (Dox) treatment for six hours. Bars represent the mean bound over input (Bd/In) normalised to the IAP gene and points represent two replicates. See Extended Data Fig 10a for ChIP-qPCR using additional antibodies. Right panel: Brightfield and GFP-fluorescence images of the mECS cells before and after Dox treatment. **h,** ATAC-seq signal reporting accessibility of the integrated locus before and after Dox treatment for six days. From left to right: annotated are the TetO array and its proximal PGK promoter that controls the Puromycin-GFP reporter gene, and the distal PGK promoter. **i,** Model: PRC1 forms multivalent interactions with chromatin, thereby stabilizing chromatin condensates potentially through charge screening of negatively charged DNA by positive charges in the CBX8 IDR. These interactions dynamically change as PRC1 diffuses through condensates.

Data thus far suggest that the deletion of CBX8 does not change chromatin accessibility in mESCs (Fig.5c,f). Yet, other CBX proteins could potentially compensate for the loss of CBX8 in endogenous polycomb target genes. Therefore, we wished to test CBX8 in a system where its recruitment is sufficient to trigger gene repression. To this end, we used a mESC line that expresses a TetR-CBX8 fusion and includes a GFP reporter cassette downstream of a TetO DNA binding array, stably integrated at chromosome 15 (coordinates: mm10 chr15: 79,013,675; Fig. 5g) ^42^. In the absence of doxycycline (Dox), the TetR-CBX8 fusion is recruited to the TetO-GFP reporter together with RING1B and initiates transcriptional repression (Fig. 5g and Extended Data Fig. 10a). However, GFP repression is reversible upon Dox addition, which causes release of both TetR-CBX8 and RING1B from the TetO DNA binding array (Fig. 5g and Extended Data Fig. 10a).

Remarkably, ATAC-seq after prolonged Dox treatment did not reveal a substantial increase in chromatin accessibility over the promoter region (Fig 5h). Small changes in the ATAC-seq signal can be measured across samples and replicates, but these follow the sample-to-sample variations globally (see Extended Data Fig. 10b for ATAC-seq coverage over the HoxA genes cluster) and are therefore unlikely to be indicative of increased local accessibility.

The only increment in accessibility upon Dox treatment occurred downstream of the GFP cassette, at a considerable distance from the TetO recruitment site and its adjacent PGK reporter (Fig. 5h). That distal site includes another PGK promoter that is placed over 5 kbp downstream to the TetO array. That distal PGK promoter originated from the construct that was used to generate the reporter (^42^ and references therein) and does not express a functional protein-coding mRNA. Hence, we cannot exclude the possibility that CBX8 affects accessibility at sites distant from the main recruitment hub, possibly through indirect effects. Nevertheless, the data indicate that CBX8 recruitment and subsequent gene repression are insufficient to restrict chromatin accessibility at the recruitment site.

## Discussion

In conclusion, we have shown that PRC1-CBX8 binds chromatin via multivalent interactions and induces chromatin condensation using both the nucleosome interacting surface of PRC1 and the IDRs of CBX8. PRC1^C8^ is dynamic within condensates while keeping chromatin in a static, solid-like state (Fig 3). This is in contrast to the liquid-like state of chromatin condensates that were observed in vitro, in the absence of PRC1^17^. We established that PRC1^C8^ is sufficient to induce the solid condensed state of chromatin.

How can PRC1 condense chromatin but yet maintain a highly dynamic behaviour in the nucleus? PRC1 is characterised with a short residence time on chromatin^31^. Solid-like chromatin that is condensed by a mobile chromatin binder has been shown *in vitro* for a truncation of the SAM-domain protein Polyhomeotic (Ph)^43^ and in cells for HP1a at chromocenters^20^. Yet, the mechanism allowing PRC1 and other gene-repressing factors to condense chromatin while constantly diffusing in it remained largely unknown.

Our cryo-EM structure of PRC1-condensed chromatin explains how PRC1 can move in condensed chromatin (Fig. 1), owing to the large pores that are formed between condensed nucleosomes. The multivalent interactions between PRC1-CBX8 to chromatin provide PRC1 with multiple docking sites on chromatin: unmodified and modified H3 tail (Fig 3b-d), DNA (Fig 3c) and the acidic patch of the nucleosome^36^. Hence, it is possible that PRC1-CBX8 can constantly change its interactions with chromatin to maintain its condensed structure while utilising its different chromatin-interacting surfaces to dynamically move around. While doing so, the positively charged IDRs of CBX8 mask the negative charge of the DNA to bring together chromatin segments and stabilize the condensed state of chromatin (Fig. 5i).

Neither the DNA binding activity of the chromo domain (Fig. 4b) nor H3K27me3 (Fig. 3d) seem to be necessary for efficient PRC1^C8^-chromatin phase separation. A potential limitation in the usage of a methyl-lysine analogue (Fig 3d) is that it may not always serve as a perfect histone mimic (discussed in ^37^). However, since the chromodomain of CBX8 is dispensable for chromatin condensation (Fig. 4), it is plausible that histone tail binding may be dispensable too. Given that the chromodomain is required for DNA binding (Fig. 4d) and implicated in binding to H3K27me3^37,38^, it may play a role predominantly in recruitment.

In cells, canonical PRC1 includes an additional PHC protein, which was previously implicated in chromatin compaction and condensation ^8,28,43^. We reasoned that an in vitro study of a simplified three-subunit complex, devoid of a PHC subunit, would allow us to characterize the chromatin condensation activity of the CBX subunit. This minimal complex also allowed us to overcome difficulties in purifying a PHC-bound PRC1 complex in sufficient quantities and purity for structural studies. It is plausible that the chromatin compaction activities of CBX8 and the PHC subunit cooperate in vivo and the absence of a PHC subunit in our experiments presents a limitation when trying to extrapolate from our in vitro results.

While our experiments were not designed to extensively characterise the contribution of protein-protein interactions towards phase separation or chromatin condensation, we cannot exclude their involvement. Indeed, oligomerization of BMI1 was reported^44^ and PHC polymerization influences PRC1-chromatin condensate properties^28^. However, PRC1^C8^ does not form condensates in the absence of chromatin in vitro (Fig 1e). Furthermore, there are only about 10 PRC1 molecules per polycomb domain in cells^31^, which is a factor that needs to be considered when attempting to link protein oligomerization to chromatin compaction. More studies are needed to determine how PRC1 molecules are distributed and work together within polycomb domains in vivo and how protein-protein and protein-DNA interactions contribute to this process.

Our data indicate that chromatin condensation together with dynamic behaviour within chromatin condensates is an intrinsic biophysical property of PRC1-CBX8. It is plausible that the dynamic behaviour of PRC1 within chromatin condensates is required in order to allow PRC1 to modify nucleosomes by the H2AK119ub mark while holding them together. This phenomenon might represent a broad paradigm of repressive chromatin. The internal structure of PRC1-chromatin condensates is a porous network of nucleosomes (Fig. 1). Such a structure could present a size-selective diffusion barrier, in agreement with its permeability to PRC1 diffusion in vitro (Fig. 2) and Tn5 accessibility in cells (Fig. 5). The existence of such a size-selective diffusion barrier remains to be identified in vivo, where it may contribute to gene repression by selectively excluding transcriptional coactivators, which are commonly large protein complexes (>1MDa^12,45–47)^. ATAC-seq experiments reach their limitations in this context, because of the small size of the Tn5 used in these assays.

Hence, from our experiments in cells, we can only conclude that CBX8-bound chromatin is not entirely inaccessible. Future studies may develop size-selective probes to directly address questions of size-selective chromatin accessibility genome-wide. The hypothesis that polycomb-mediated repression antagonises Pol II transcription without blocking all proteins has been made nearly three decades ago^48^. This idea was conceived based on the observation that T7 polymerase (∼100 kDa) can initiate transcription from a polycomb-repressed locus but GAL4-dependent transcriptional activation does not take place there. This is in agreement with the inverse correlation between the density of chromatin domains and the molecular weight of the chromatin modifiers present in them^12^. Transcription factor size has also been suggested to determine access to different chromatin domains based on simulations^49^. Combining our results with earlier findings^12,19,20,49^, we propose that size-selective exclusion may be part of a broader mechanism by which chromatin-interacting proteins regulate the accessibility of repressive chromatin.

## Acknowledgements

We would like to thank the Ramaciotti centre for electron microscopy at Monash university for providing instrumentation, technical support and collecting cryo-ET data, the Monash Proteomics facility for providing instrumentation and technical support and the Monash Micro Imaging facility for providing instrumentation and technical support. We would also like to acknowledge the MASSIVE HPC platform for providing high-performance computing resources

## Authors contributions

C.D. and M.U. conceptualised the project and acquired funding, M.U., V.L., C.T., X.H.N, J.M., J.H., S.M., H.V., S.T., M.B. and M.L. carried out experiments and investigated, P.P.D. and V.P. generated cell lines, S.F. and Q.Z. cloned and purified histone mutants, M.L. and M.U. developed software, C.D., A.d.M., O.B., P.P.D., Y.G. and E.G. supervised, M.U. and C.D. wrote the original draft and all authors reviewed and edited the manuscript.

## Data and materials availability

Next generation sequencing data (ATAC-seq and ChIP-seq) are available under GEO accession number GSE220140. The maps for tomograms in Fig. 1 g, and j have been deposited to the EMD (EMD-29022 and EMD-43554, respectively). All cryo-ET raw data has been deposited to EMPIAR under accession numbers EMPIAR-11344 and EMPIAR-11883, of chromatin in the presence and absence of PRC1-CBX8, respectively. XL-MS data has been deposited to Pride (PXD039589 and PXD049094).

## Extended Data

**Extended Data Fig. 1.**
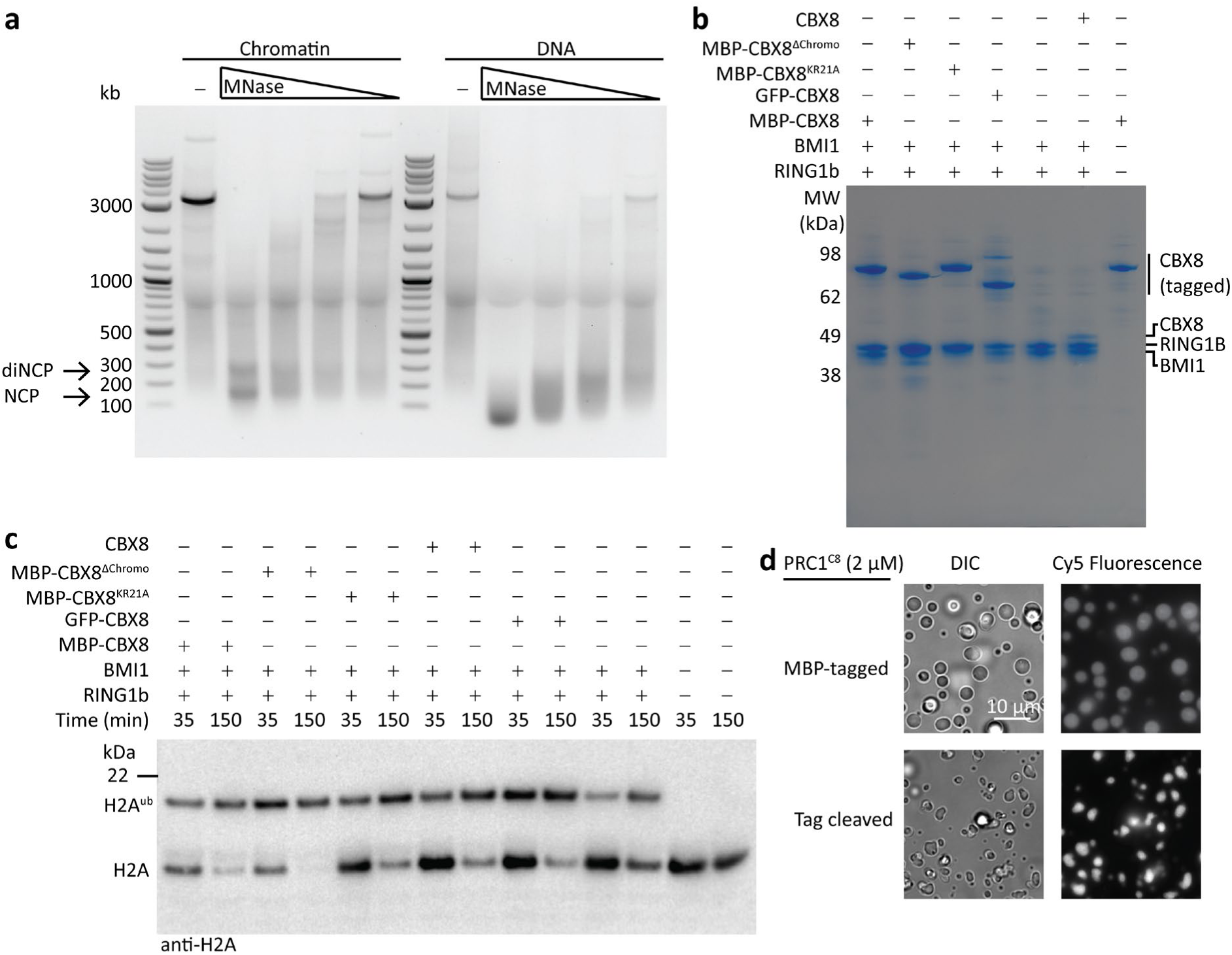
Quality control of chromatin and protein constructs. **a,** MNase digestion of reconstituted Chromatin and a naked DNA (same sequence as used for chromatin reconstitution). DNA fragments post digestion are resolved on a 1.2 % Agarose gel. Protected bands indicating mono- and di-nucleosome core particles (NCP and diNCP) are indicated by the arrows. **b,** 3 ug of each protein complex used in this study resolved on a 4-12% SDS-PAGE gel stained with Coomassie. **c,** Ubiquitylation activity of each protein complex used in this study visualized on a western blot. All samples include UBA1, UBCH5C, Ubiquitin, ATP and 1 uM chromatin (nucleosome concentration). **d,** Phase separation experiment comparing chromatin condensation activity of PRC1^C8^ with MBP-tagged CBX8 to PRC1^C8^ with the tag cleaved.

**Extended Data Fig. 2.**
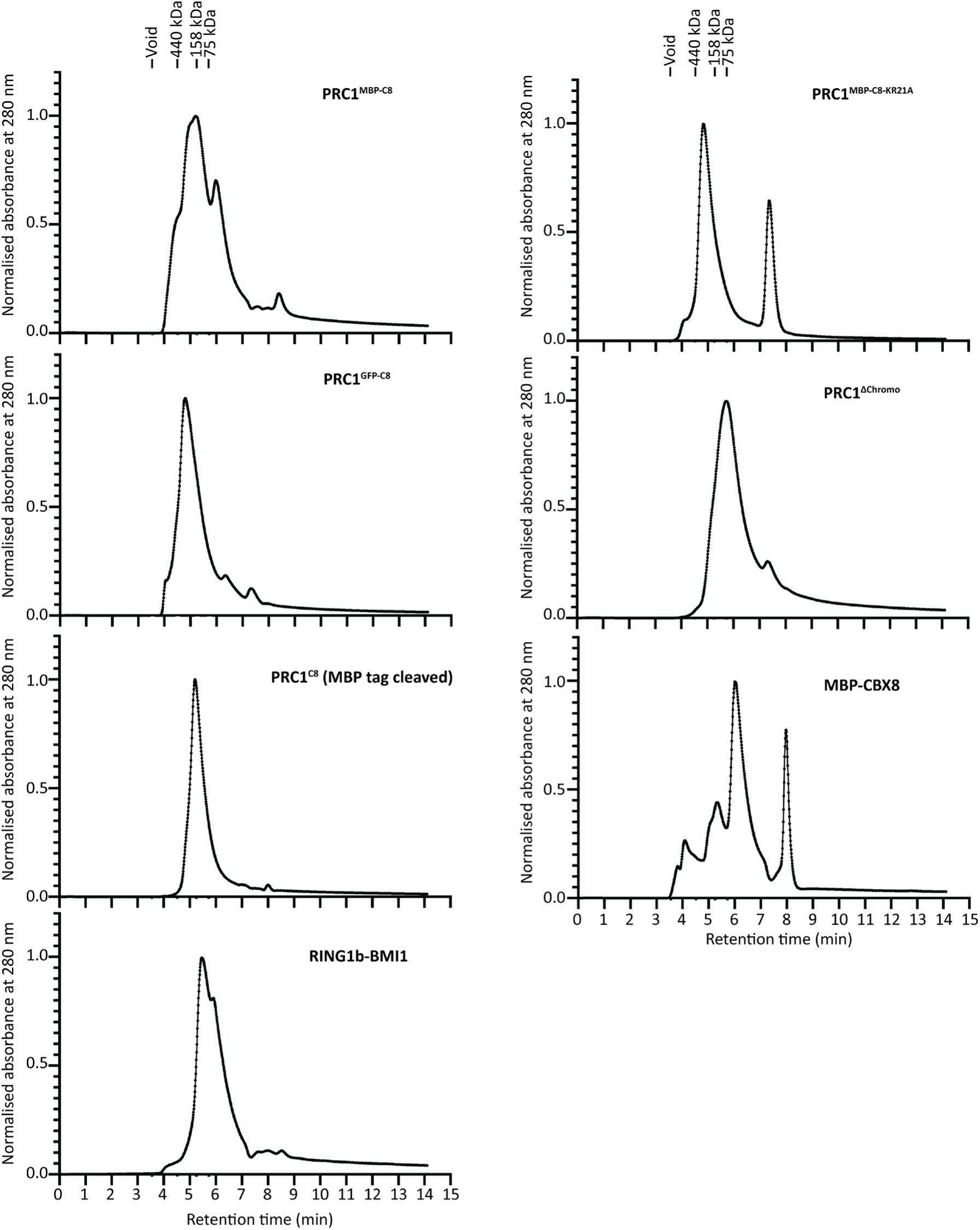
PRC1 complexes used in this study are monodispersed. HPLC elution profiles from a Shim-pack Bio Diol 200 HPLC column for each of the purified protein complexes, as indicated.

**Extended Data Fig. 3.**
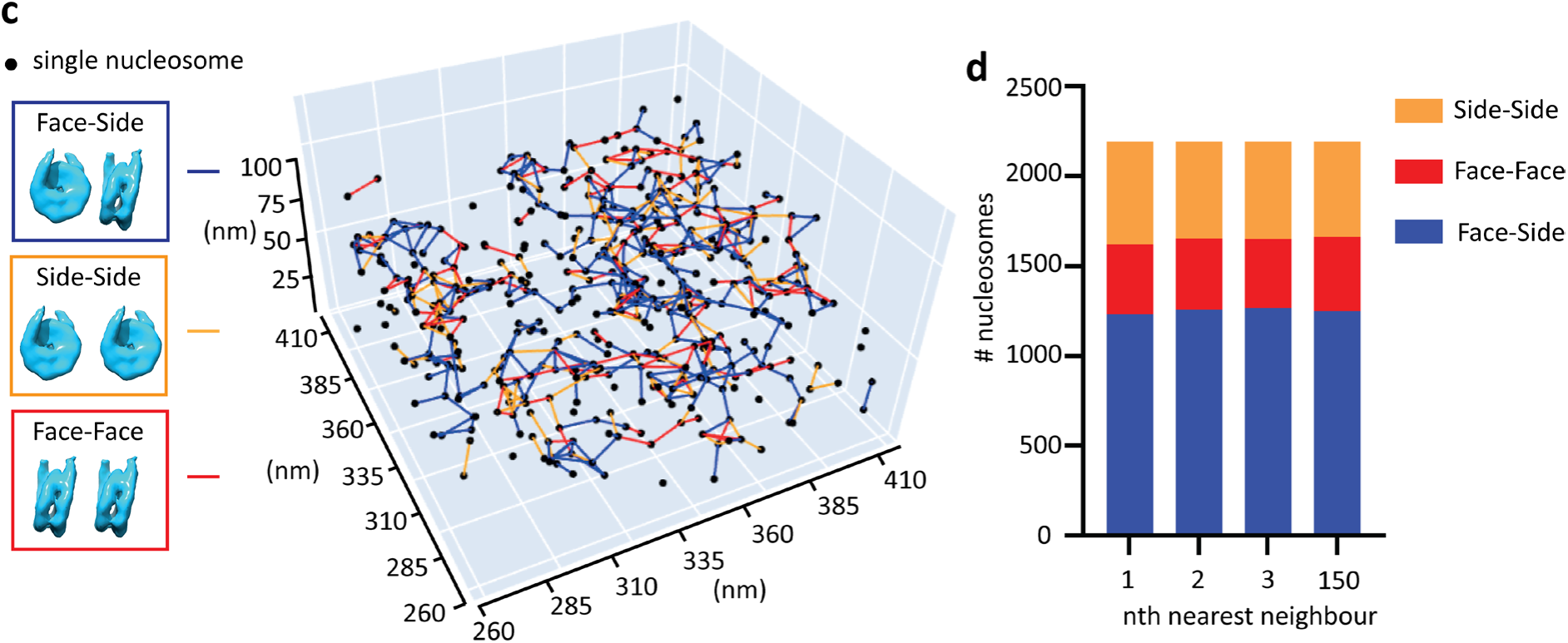
Nucleosomes in PRC1^C8^-chromatin condensates show no orientation bias towards neighbouring nucleosomes. **a,** Low-magnification micrograph showing examples of dense regions suspected to be condensates (dashed circles). Tomograms were collected at the borders of regions such as these. Micrographs are from the same gird from which tomograms were collected. **b,** Structures of the template used for template matching (left), the averaged structure after template matching (middle) and the final structure after the subtomogram averaging routine (right) with the related Fourier shell correlation curve (bottom). **c,** Orientations of nucleosomes towards neighbouring nucleosomes within a cut-off of 20 nm. Individual points represent nucleosomes and lines between points are coloured according to the relative orientation of neighbouring nucleosomes as indicated in the colour key (left). **d,** Distribution of nucleosome-nucleosome orientation for the three nearest neighbours and the 150th neighbour of each nucleosome in tomogram #1. Colours correspond to the same respective orientations as in **c**.

**Extended Data Fig. 4.**
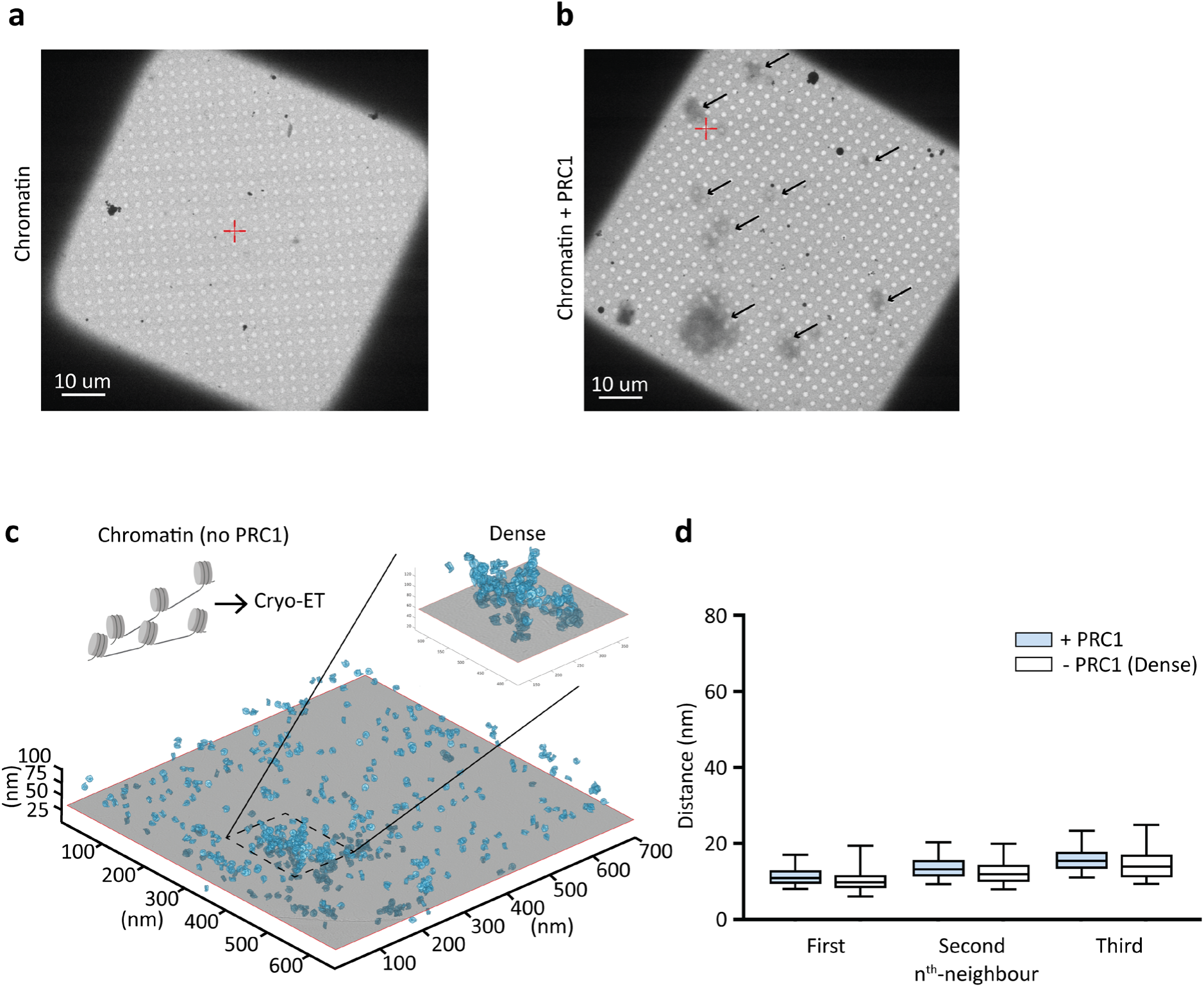
**a,** Representative low-magnification cryo-EM micrograph of a grid square with vitrified chromatin in absence of PRC1. **b,** Same as (**a**) but with PRC1. Black arrows indicate presumed condensates. Red crosses relate to stage movement and do not indicate features in the context of this figure. **c,** The same cryo-tomogram as in Fig. 1j, of chromatin in the absence of PRC1, with dense regions highlighted. **d,** Distances to the three nearest neighbouring nucleosomes, where +PRC1 as in Fig. 1k and -PRC1 shows distances for the highlighted dense region in **c**.

**Extended Data Fig. 5.**
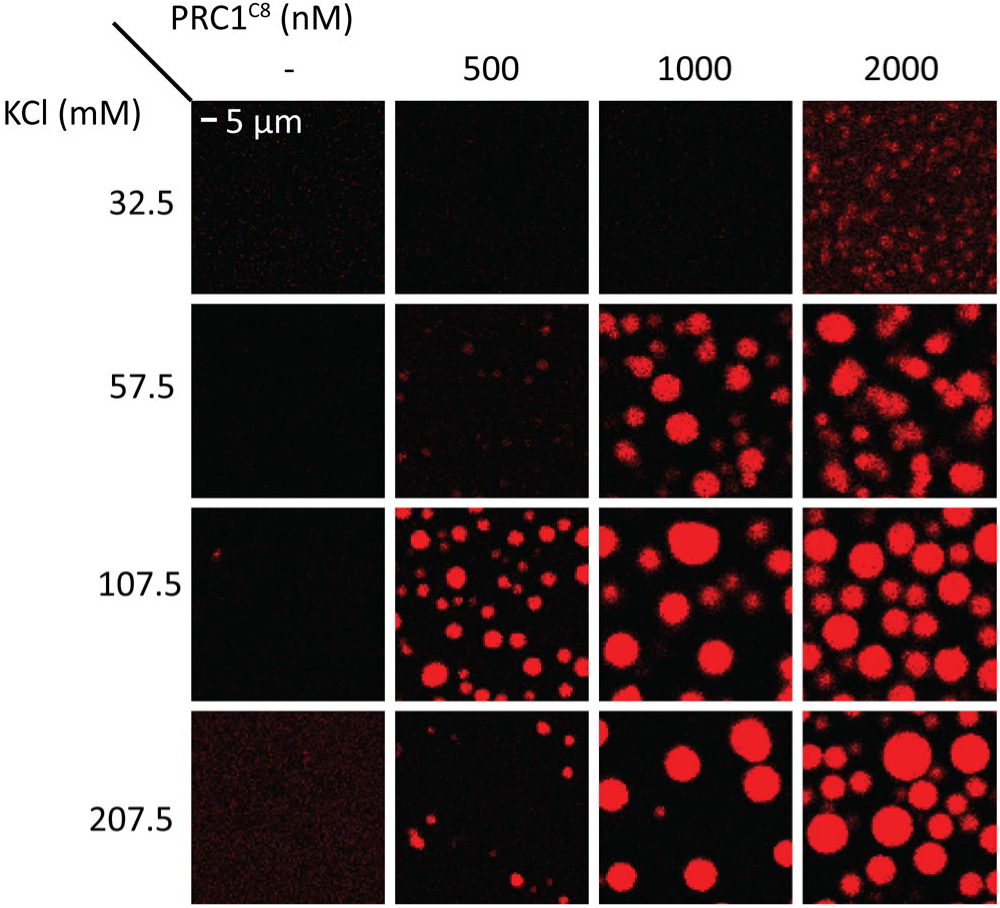
Chromatin condensation in response to variations in salt and PRC1 concentration measured by confocal microscopy using Cy5-labelled chromatin at a constant concentration of 50 ng/μl DNA (approximately 400 nM nucleosomes).

**Extended Data Fig. 6.**
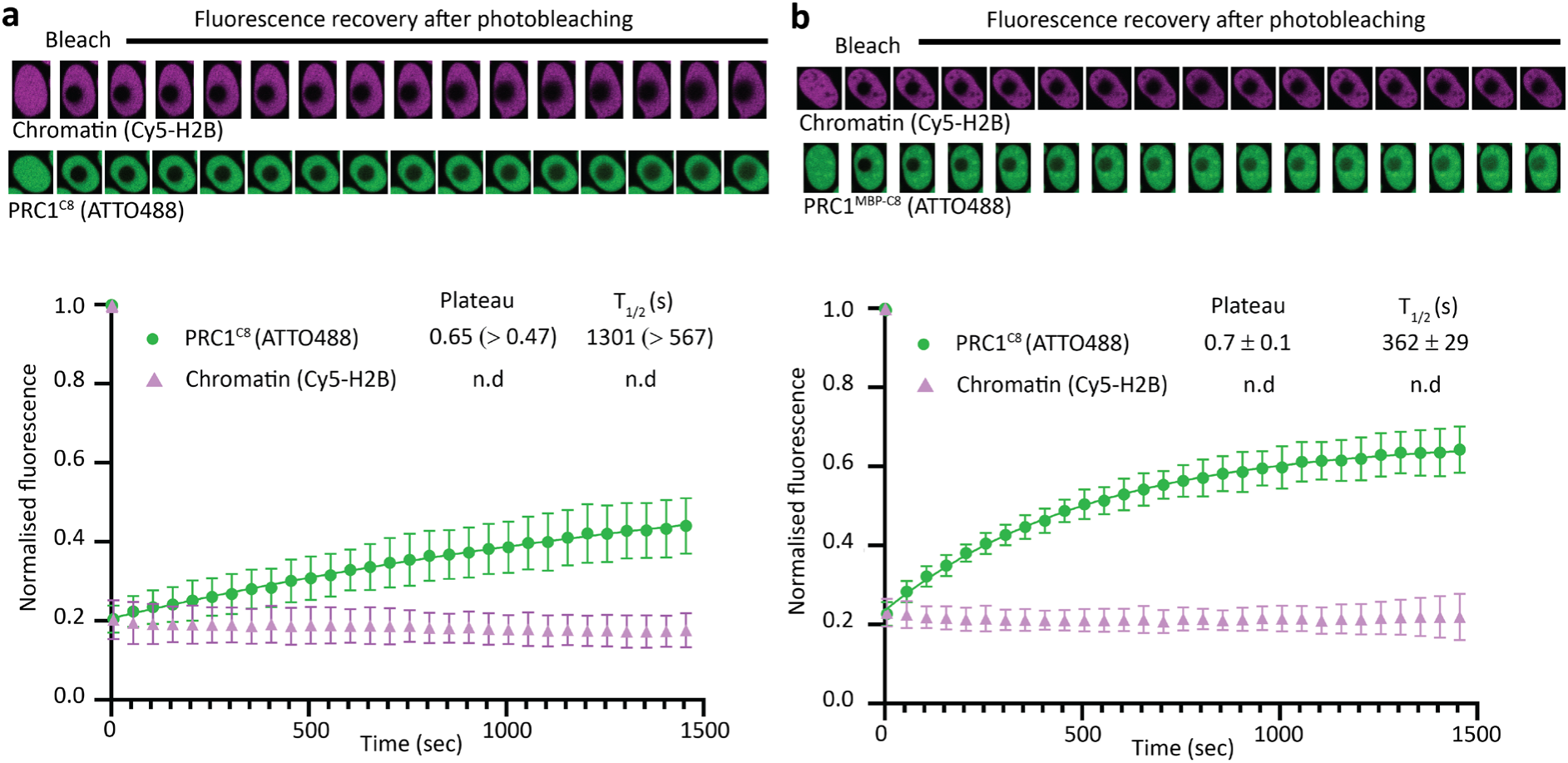
Untagged PRC1^C8^ is mobile while chromatin is static within PRC1-chromatin condensates. **a,** Representative micrographs of FRAP recorded in PRC1^C8^-chromatin condensates, where within this complex CBX8 is untagged. PRC1^C8^ is labelled with ATTO488 and chromatin is Cy5 labelled. The mean fluorescence intensity of the bleached area, normalised to the pre-bleach mean signal, is plotted for every time point. Error bars show standard deviation from n=6 (GFP) and n=7 (Cy5) measurements that were recorded from four independent experiments that were carried out on different days. The GFP signal recovery was fit with an exponential association model, best fit values for Plateau and fluorescence recovery half time (T_1/2_) are shown. The lower limits of the 95% confidence interval are presented in parentheses (the upper boundaries could not be determined confidently). **b,** Representative micrographs of FRAP recorded in PRC1^C8^-chromatin condensates, where within this complex CBX8 includes an N-terminal MBP-tag. PRC1^C8^ is labelled with ATTO488 and chromatin is Cy5 labelled. The mean fluorescence intensity of the bleached area, normalised to the pre-bleach mean signal, is plotted for every time point. Error bars show standard deviation from n=7 (GFP) and n=6 (Cy5) measurements recorded from four independent experiments that were carried out on different days. The GFP signal recovery was fit with an exponential association model, best fit values for Plateau and fluorescence recovery half time (T_1/2_) are shown with SEM.

**Extended Data Fig. 7.**
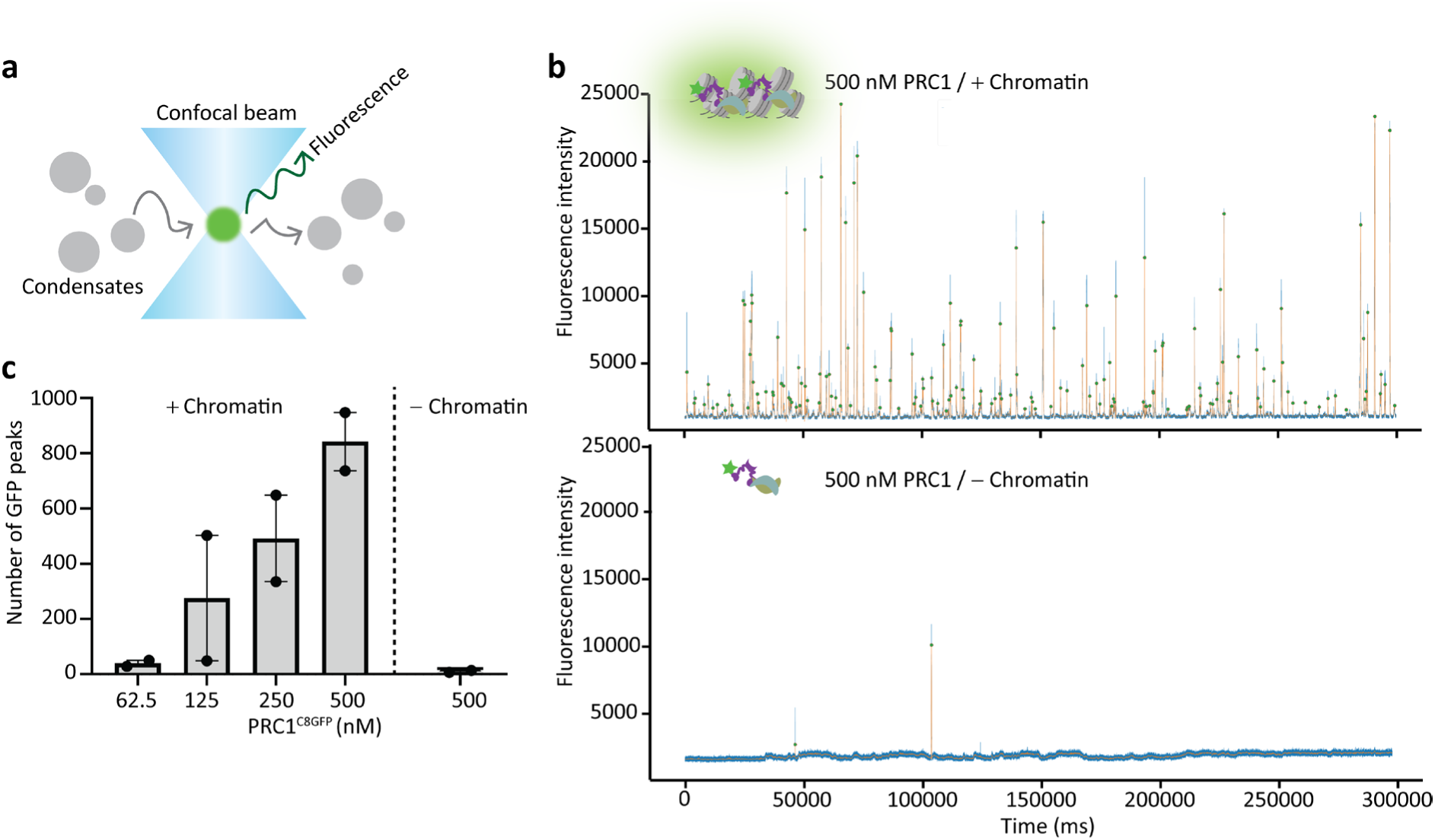
Condensates can be detected at physiologically relevant PRC1^C8^ concentrations. **a,** An illustration of single molecule confocal microscopy. Individual condensates (in grey) are detected when diffusing through the confocal volume (in blue) and emitting GFP fluorescence (in green). **b,** Top: representative trace tracking GFP signal over time for samples with 500 nM GFP-labelled PRC1^C8^ and chromatin (200 nM nucleosome concentration). Traces show a 5 minute window from a 20 minute experiment. Blue lines show the raw GFP signal and orange lines show the GFP signal after low-pass filtering using a Butterworth filter. Green dots indicate the maxima of the detected peaks. Bottom: same as the top plot, but in the absence of chromatin. **c,** GFP peak counts at different PRC1^C8^ concentrations. Data from two replicates are shown. Bars indicate the mean of two independent replicates that were carried out on different days, the error bars show the standard error of the mean and individual data points are presented.

**Extended Data Fig. 8.**
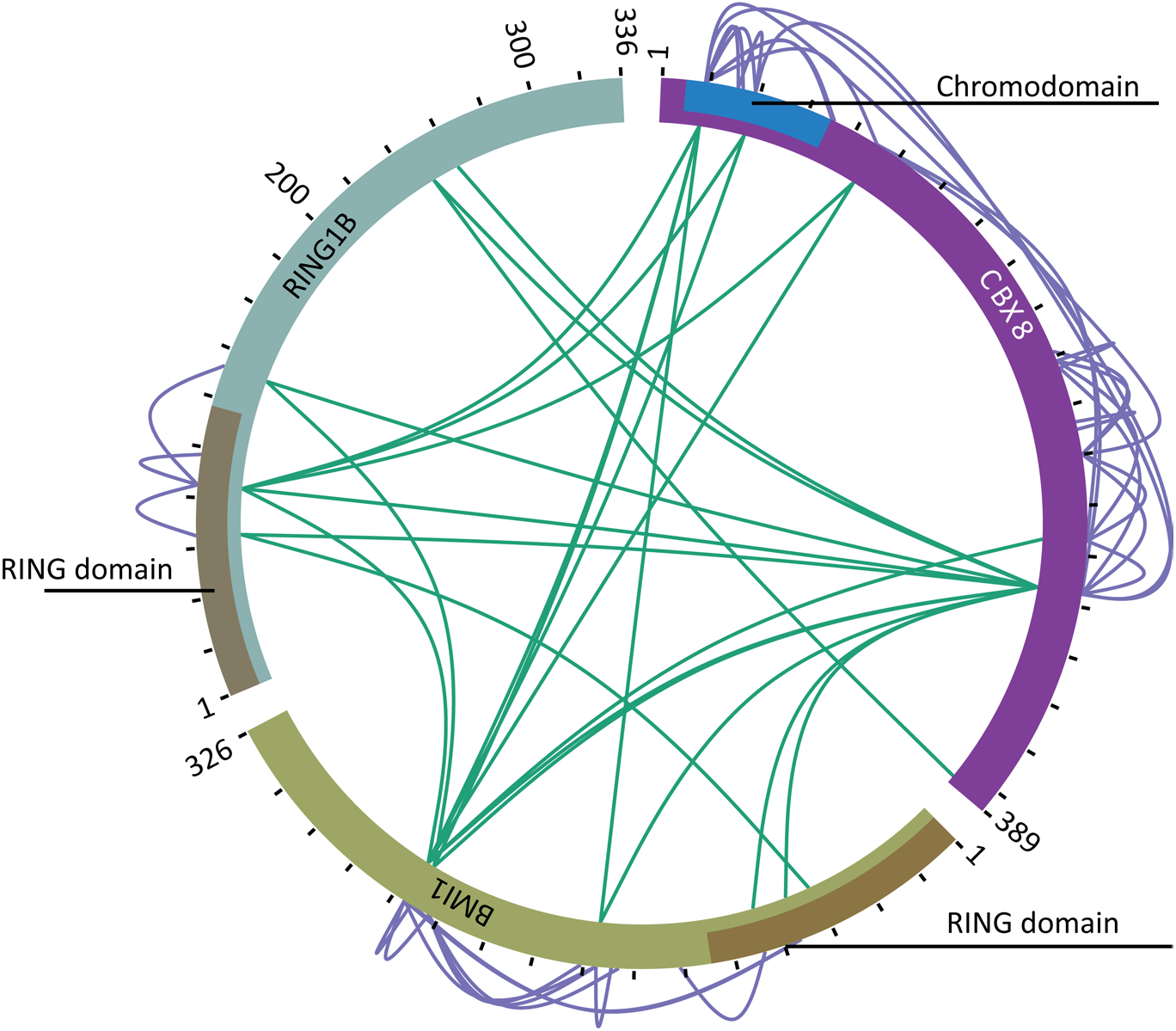
Crosslinking mass spectrometry (XL-MS) profile of PRC1^C8^ is similar in the absence of chromatin. Same experiment as in Fig. 3b, but without chromatin.

**Extended Data Fig. 9.**
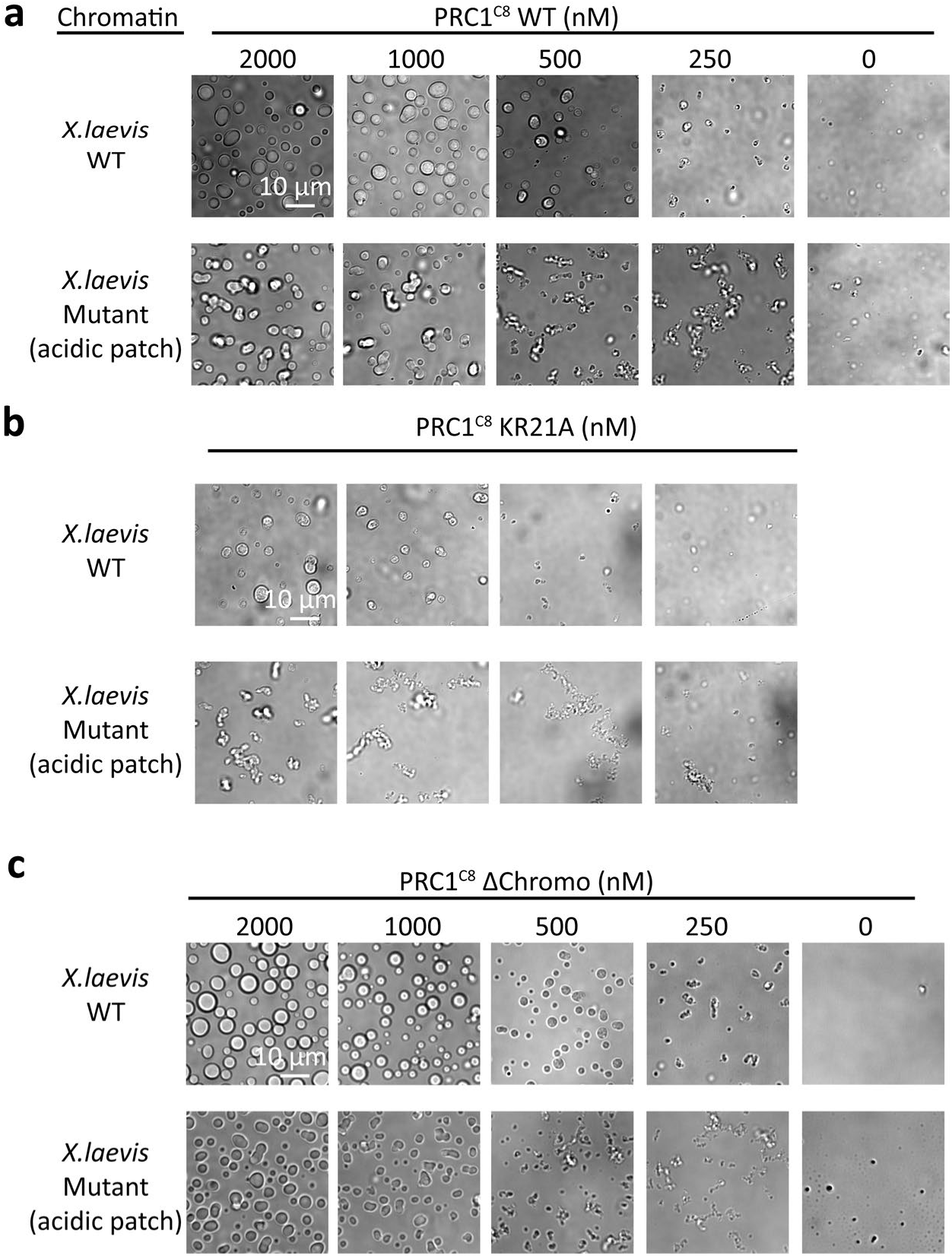
The nucleosome acidic patch is required for efficient PRC1-chromatin phase separation. **a** Phase separation of *X. laevis* wildtype and acidic patch mutant chromatin in response to increasing concentrations of PRC1^C8^ wildtype. DIC micrographs are representative of two replicates. **b** Same as (**a),** but with the PRC1^C8KR21A^ mutant. **c** Same as (**a)** but with the PRC1^C8ΔChromo^ mutant.

**Extended Data Fig. 10.**
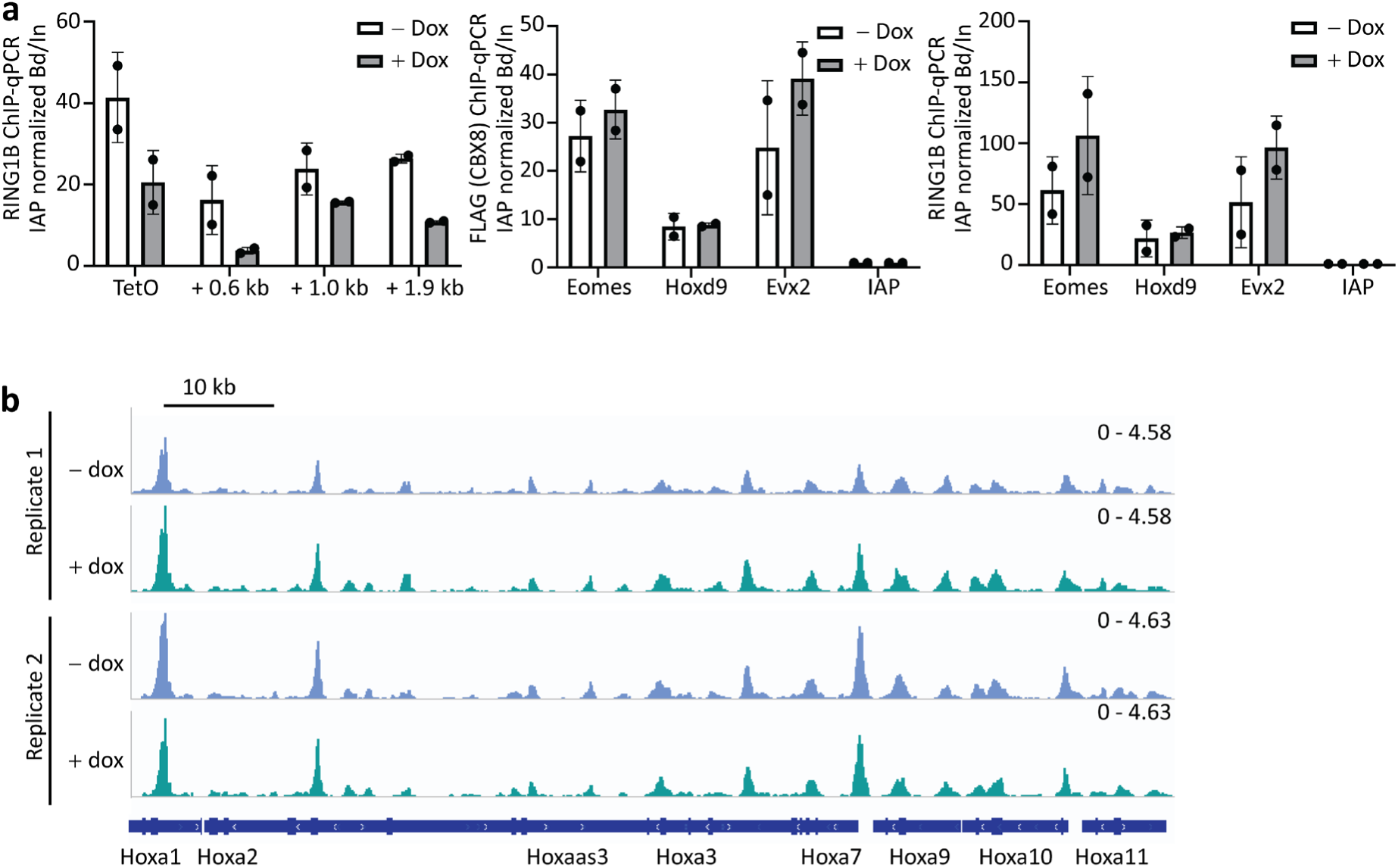
**a,** ChIP-qPCR in the presence and absence of doxycycline (Dox) treatment using RING1B antibody at indicated distances from a chromosome-integrated TetO array (left), at control genes (right) and using FLAG (CBX8) antibodies at control genes (middle). Bars represented the bound over input (Bd/In) normalised to the IAP gene and the dots represent individual data points. **b,** ATAC-seq signal at the HoxA locus of the reporter-integrated mECS cell line before and after dox treatment. Two independent replicates are shown.s

**Extended Data Movie S1. Related to Fig.1; Cryo-tomogram of a PRC1^C8^-chromatin condensate.** The movie shows a scan through the z-axis of denoised^50^ tomogram of chromatin in presence of PRC1.

**Extended Data Movie S2. Related to Fig.1; Cryo-tomogram of chromatin without PRC1.** The movie shows a scan through the z-axis of denoised^50^ tomogram of chromatin in absence of PRC1.

**Supplemental Table 1. Template matching results for PRC1^C8^-chromatin condensates.** Results from the Dynamo template matching process before manually removing false positive hits, in Excel format. Every entry describes a cross correlation peak above the cut-off of 0.17. Each peak indicates the position of a nucleosome at x-y-z coordinates given in columns 24-26 and at rotations applied to the template as defined by Euler angles in columns 7-9. The table is also provided in Dynamo format (.tbl) for direct import into Dynamo (Supplementary Data 1).

**Supplemental Table 2. Template matching results for chromatin without PRC1^C8^.** Results from the dynamo template matching process before manually removing false positive hits in Excel format. Every entry describes a cross correlation peak above the cut-off of 0.23. Each peak indicates the position of a nucleosome at x-y-z coordinates given in columns 24-26 and at rotations applied to the template as defined by Euler angles in columns 7-9. The table is also provided in Dynamo format (.tbl) for direct import into Dynamo.

**Supplementary Data 1. Template matching results for chromatin without PRC1^C8^.** Same as Supplemental Table 1, provided in Dynamo format (.tbl) for direct import into Dynamo.

**Supplementary Data 2. Template matching results for PRC1^C8^-chromatin condensates.** Same as Supplemental Table 3, provided in Dynamo format (.tbl) for direct import into Dynamo.

**Supplementary Data 3. XL-MS-identified crosslinks within PRC1^C8^.** Provided is a list of the crosslinks that were detected using XL-MS within PRC1^C8^ in the absence of chromatin.

**Supplementary Data 4. XL-MS-identified crosslinks within PRC1^C8^-chromatin.** Provided is a list of the crosslinks that were detected using XL-MS within PRC1^C8^ in the presence of chromatin.

## Materials and Methods

### Plasmids and cloning

Human RING1b (Uniprot ID Q99496) and human BMI1 (Uniprot ID P35226) were cloned into a pFBOH-mhl vector (Addgene plasmid # 62304) cleaved with BseRI using Gibson Assembly® Master Mix (NEB #E2611L) using the primers indicated in Table S1.

Human CBX8 wildtype open reading frame (Uniprot ID Q9HC52-1 and NCBI Reference Sequence was NM_020649.2) was obtained from gene synthesis (Gen9). The CBX8^KR21A^ mutant open reading frame was codon optimised for expression in *Trichoplusia ni* insect cells and synthesised (Genscript). The CBX8^ΔChromo^ truncation was amplified from the wildtype ORF using primers indicated in Table 1 and then subcloned into a vector with the same backbone as used to expresses the wild type protein. All three CBX8 constructs were cloned into a modified pFastBac1 pFB1.HMBP.A3.PrS.ybbR vector digested by XhoI and XmaI sites to include a N-terminal 6xHis-MBP tag Cloning of the polycomb target gene ATOH1 into the pUC18 vector was described previously^51^. Plasmids for expression of human histones (H2A, H2B, H3.1 and H4) in *E.coli* were a kind gift from David Tremethick, Australian National University. UbcH5c WT pET28a was a gift from Rachel Klevit (Addgene plasmid # 12643; http://n2t.net/addgene:12643; RRID:Addgene_12643)^52^. pET3a-hUBA1 was a gift from Titia Sixma (Addgene plasmid # 63571; http://n2t.net/addgene:63571; RRID:Addgene_63571)^53^. To generate a baculovirus expression vector of a monomeric EGFP-CBX8 (mEGFP-CBX8) construct, first GFP was amplified from a pSpCas9(BB)-2A-GFP vector and CBX8 was amplified from a pFB1.HMBP.A3.PrS.ybbR vector containing CBX8 as an insert. Subsequently, EGFP-CBX8 was cloned into the pFBOH-mhl vector cleaved with BseRI via Gibson assembly, with a Serine-Glycine-Serine linker between EGFP and CBX8. Finally, to generate monomeric mEGFP-CBX8, alanine residue 207 in EGFP was mutated to Lysine using a site directed mutagenesis kit (Takeda) and the mutagenesis primers listed in Table 1.

**Table 1:**
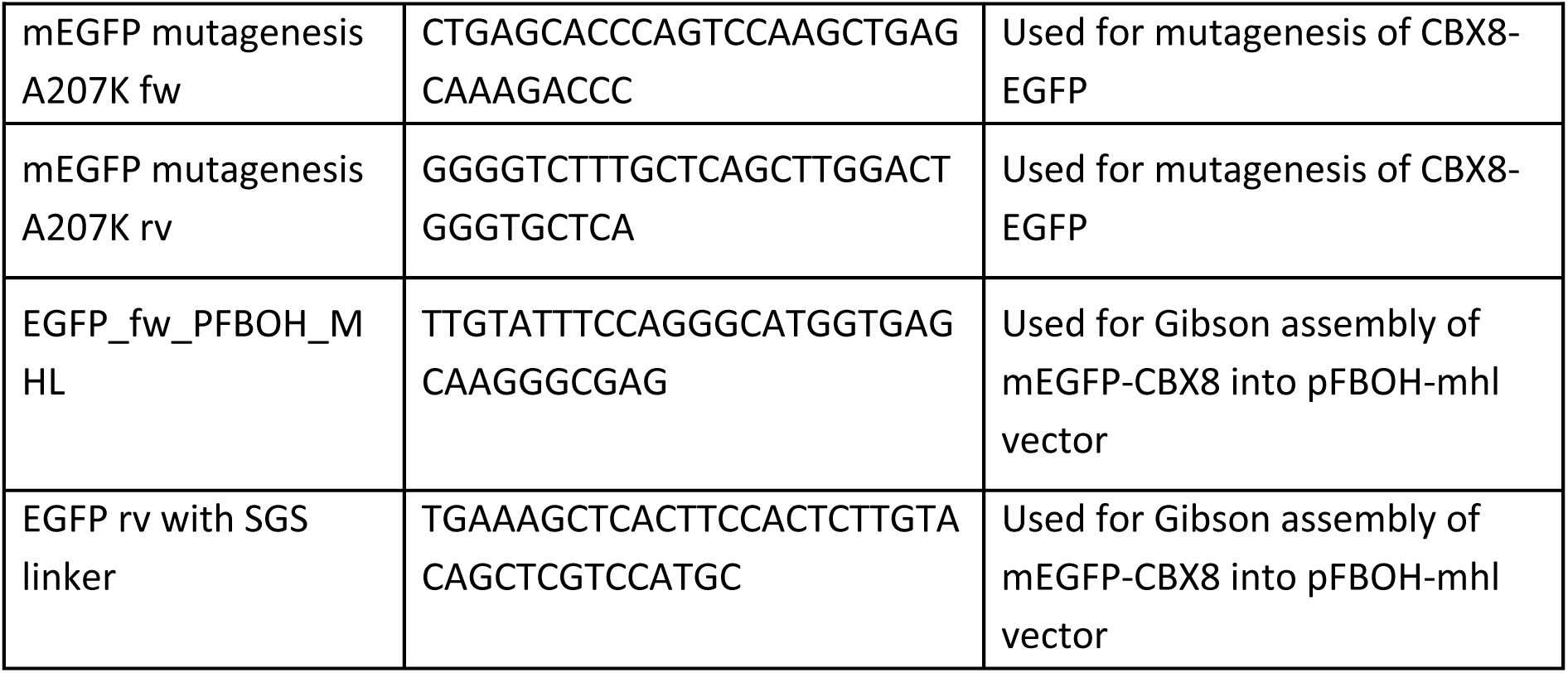
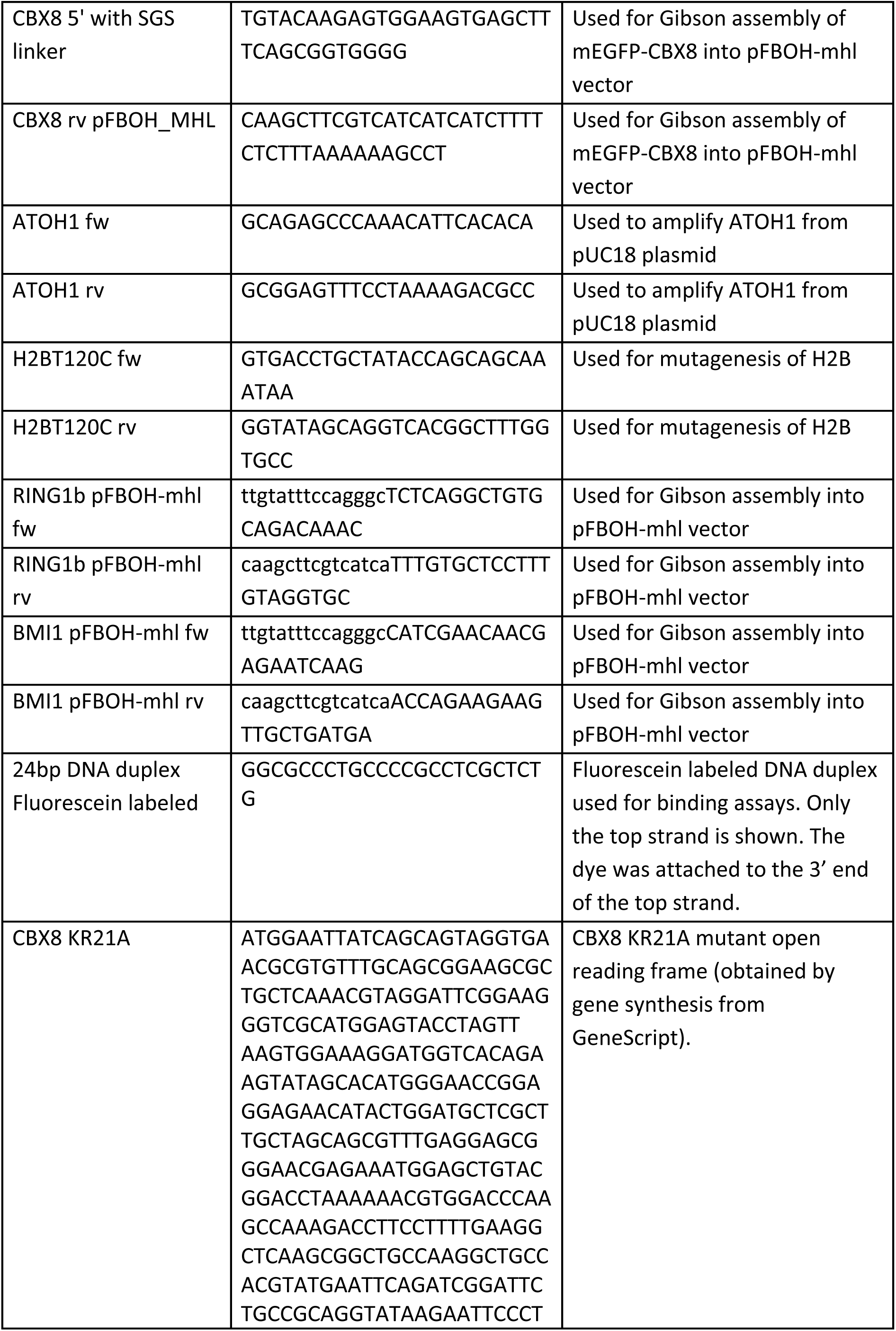
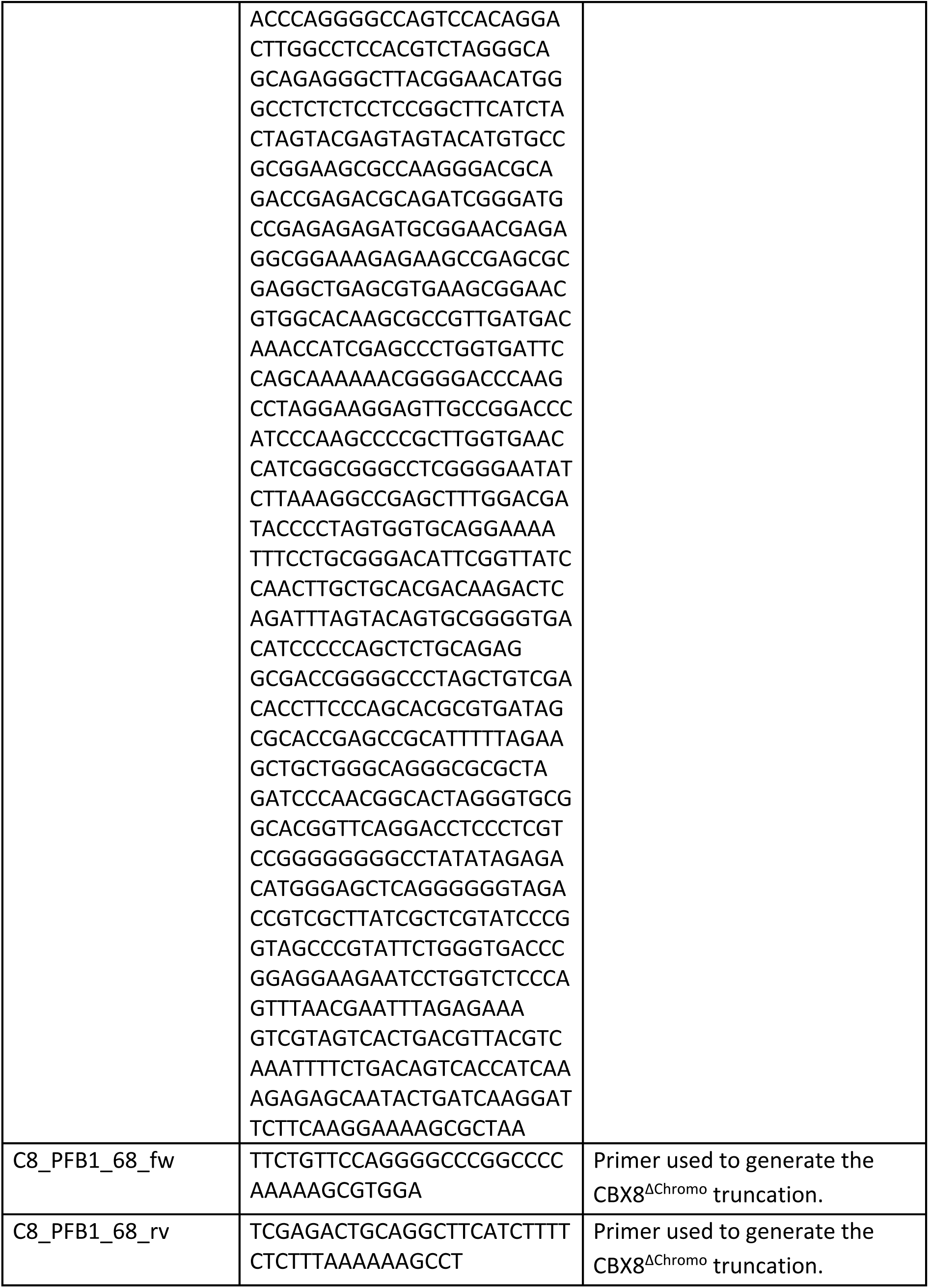
Cloning and mutagenesis primers (5’-3’) and sequences.

### Protein expression and purification

PRC1^C8^, PRC1 core and CBX8 were co-expressed in *Trichoplusia ni* insect cells using the Baculovirus system. CBX8 variably carried an N-terminal 6xHis-mEGFP or a N-terminal 6xHis-MBP tag. The purification protocols were identical regardless of the tag. Baculoviruses were generated as per manufacturers instructions (Thermo Fisher). The viral titre was determined using the MTT assay (Promega #G3580). *Trichoplusia ni* insect cells were infected at a density of 1.5 - 2 × 10^6^ cells and incubated for 60 hours at 27 °C in a shaker. The cells were spun down at 1500 relative centrifugal force (RCF). The pellet was resuspended in 100 ml of lysis buffer per litre of cell culture (20 mM HEPES-KOH pH 7.5, 400 mM NaCl, 25 mM Imidazol, 10 % Glycerol, 0.2 mM TCEP, 1 mM PMSF and EDTA-free Complete protease inhibitor (Thermo Fisher)). Lysis proceeded for 30-45 minutes at 4 °C while rotating. The lysate was then centrifuged at 29000 RCF for 20 minutes at 4 °C. The supernatant was transferred to a fresh tube and 5 ml of Ni-NTA resin (Qiagen) was added. The samples were then incubated for 60 minutes at 4 °C while rotating and subsequently centrifuged at 500 RCF for 5 minutes at 4 °C to settle the beads. About 90 % of the supernatant was removed. The beads were resuspended in the remaining 10 % of supernatant and transferred to 25 mm diameter gravity flow columns (Biorad). The beads were allowed to settle before the remaining buffer was drained and the beads were then washed with 12 CV of Buffer B (20 mM HEPES-KOH pH 7.5 at 20 °C, 500 mM NaCl, 25 mM Imidazole, 10 % Glycerol, 0.2mM TCEP), followed by 30 CV of Buffer A (20 mM HEPES pH 7.5 at 20 °C, 100 mM NaCl, 25 mM Imidazole, 10 % Glycerol, 0.5 mM DTT or 0.2 mM TCEP). The protein was then eluted in 6 CV Elution Buffer (20 mM HEPES-KOH pH 7.5 at 20 °C, 100 mM NaCl, 400 mM Imidazole, 10 % Glycerol, 0.2 mM TCEP). The eluted protein was loaded onto a Hitrap 5 ml Heparin column (Cytvia) equilibrated in IX Buffer A (20 mM HEPES pH 7.5 at 20 °C, 100 mM NaCl, 0.5 mM DDT or 0.2 mM TCEP) and the column was washed with 5 CV of IX Buffer A. The proteins were resolved over a 20 CV gradient ranging from 0 % to 100 % IX Buffer B (20 mM HEPES pH7.5 at 20 °C, 1000 mM NaCl, 0.5 mM DTT or 0.2 mM TCEP). The fractions were analysed on SDS-PAGE and fractions containing the protein complex of interest with the expected subunits stoichiometry were pooled. The pooled fractions were concentrated using a Amicon ultra 30K centrifugal filter (Merck, cat UFC903024) and purified by size exclusion chromatography using a HiLoad Sephacryl 300 16/60 column (Cytiva) equilibrated in GF Buffer (20 mM HEPES-KOH pH 7.5 at 20 °C, 150 mM NaCl, 0.5 mM DTT). The collected fractions were analysed on SDS-PAGE and fractions containing the protein complex of interest at the expected stoichiometry were pooled and concentrated to a concentration of 1-2 mg/ml using an Amicon ultra 30K centrifugal filter (Merck, cat UFC903024). The purified protein was then aliquoted and frozen in liquid nitrogen. The purified proteins were stored at -80 °C until use.

For the production of Atto-488 labelled PRC1^C8^ complexes, purification was carried out as above, with some modifications. PRC1^C8^ was purified as described above, without cleaving the N-terminal MBP tag from CBX8, up until the end of the Ion Exchange Chromatography (HiTrap 5 ml Heparin column, as above). Then, fractions containing PRC1^C8^ were pooled together and concentrated to 2.4 mg/ml. To generate untagged PRC1^C8^, PreScission protease was added to 1:50 protease:PRC1^C8^ mass ratio in a 3.5 ml reaction volume, incubated overnight at 4-8 °C and then the MBP tag was removed using 0.8 ml amylose beads through a batch removal, before proceeding to the subsequent labelling reaction. For the fluorescence labelling of MBP-tagged PRC1^C8^, the subsequent labelling reaction was carried out without tag cleavage. For fluorescence labelling, of either MBP tagged or untagged PRC1^C8^, 2.4 mg/ml protein was supplemented with Atto-488 NHS ester (Merck 41698; the Atto-488 NHS ester was dissolved in DMSO to 4.3 mM before used) to a final molar stoichiometry of 1.4:1.0 dye:protein and allowed to incubate 1 hour at room temperature in the dark, and then the protein was loaded on a light-protected HiLoad Sephacryl 300 16/60 column equilibrated with 25 mM HEPES-KOH pH 7.5, 0.5 mM DTT, and either 150 mM NaCl or 500 mM NaCl for the MBP-tagged or untagged PRC1^C8^, respectively. The process was completed as described above, with the exception that the fluorescently labelled proteins were protected from light until experimentation.

Human UBA1, UBCH5C and Ubiquitin were purified as described previously^54^. Human histone proteins H2A, H2B, H2BT120C, H3 and H4 were purified as described previously^55^, except that the gel filtration step was omitted. Purified *Xenopus laevis* histone proteins H2A, H2B, H3, H4 and H2A acidic patch mutant (E56T, E61T, E64T, D90S, E91T, E92T) were bought form from the Histone Source of the Colorado State University, Fort Collins.

### Production and purification of DNA for chromatin reconstitution

The ATOH1 polycomb target gene was amplified in a large scale 10 ml PCR reaction including 500 nM ATOH1 fwd and reverse primers (see Table 1), 4 μg of ATOH1-pUC18 template^51^, 200 μM dNTP mix (Invitrogen), 3 % DMSO, 50 mM KCl, 10 mM Tris-HCl, pH 8.8, 1.5 mM MgCl2, 0.01% sterile gelatin. The reaction mixture was divided into 96-well plates to include 50 μl per well and the following PCR program was run:

| Temperature (°C) | Time (minutes) | Number of Cycles |
| --- | --- | --- |
| 95 | 3:00 | 1 |
| 95 | 0:15 | 25 |
| 63 | 0:20 |  |
| 72 | 8:00 |  |
| 72 | 3:00 | 1 |

The PCR product was purified via ion exchange chromatography using a 5 ml HiTrap Q column (GE Healthcare). The column was equilibrated in Buffer A (25 mM HEPES 7.5, 250 mM NaCl) and the sample was resolved using a linear gradient ranging from 0 % to 100 % Buffer B (25 mM HEPES pH 7.5, 2 M NaCl). The fractions containing pure DNA were pooled, subjected to ethanol-precipitation and resuspended in 10 mM Tris-HCl pH 7.5 at 20 °C, 0.1 mM EDTA.

### Chromatin reconstitution

Histone Octamers were refolded as described previously^55^. All steps were done at 4 °C or on ice. Chromatin was reconstituted following the salt-gradient dialysis protocol described previously^55^. For large scale reconstitution, DNA and octamer were combined at an optimal ratio that was determined at trial experiments for each batch of octamers and DNA. To determine the optimal ratio of histone octamer to DNA, titration was carried out using increasing amounts of octamer to a constant amount of DNA (molar ratios of 16:1, 18:1, 20:1, 22:1 of Octamer:DNA) following by salt gradient dialysis. For the salt gradient dialysis, samples were initially dialysed in a buffer containing 10 mM Tris-HCl pH 7.5 at 20 °C, 2 M KCl, 1 mM EDTA, 1 mM DTT. The initial buffer was then gradually exchanged for a low salt buffer containing 10 mM Tris-HCl pH 7.5 at 20 °C, 250 mM KCl, 1 mM EDTA, 1 mM DTT over the course of 18 hours, after which the salt exchange was complete. Samples were then centrifuged at 21000 RCF for one minute, separated on a 1% agarose gel in TAE buffer and stained with SYBR Safe (Sigma). The highest Octamer:DNA ratio at which chromatin remained soluble after finishing salt dialysis was finally used for large scale reconstitution. For large scale reconstitutions, the salt gradient dialysis was performed as above while scaling up the reaction accordingly. Additionally, after conclusion of the salt gradient, the samples were transferred to 400 ml of low salt buffer (10 mM TrisRIS-HCl pH 7.5 at 20 °C, 250 mM KCl, 1 mM EDTA, 1 mM DTT) and dialysed for another 2 hours. The samples were finally dialysed against 1 litre of Chromatin Storage Buffer (10 mM TRIS-HCl pH 7.5, 10 mM KCl) overnight.

### Reconstitution of Fluorescently labelled chromatin

To allow site-specific labelling of histone H2B, a cysteine was introduced via site-directed mutagenesis (H2BK120C, as previously described^56^) using mutagenesis primers indicated in Table S1. H2BK120C was labelled with Cyanine5-maleimide (Lumiprobe cat #13080) as described previously^56^. Labelled octamers were refolded as described above. Before chromatin reconstitution, the unlabelled and labelled octamers were combined at a molar ratio of 7:1 (unlabeled:labelled). Chromatin was then reconstituted as described above.

### Chromatin condensation assays

Fluorescently labelled proteins and chromatin were protected from light whenever possible. Chromatin condensation assays were done as described previously^17^ with some modifications. Assays were done in 384-well plates with #1.5 glass bottoms (MatTek PBK384G-1.5-C). The wells were treated with 1 M NaOH for 1 hour at room temperature, NaOH was removed and wells were washed with copious amounts of MilliQ water (MQ). MQ was removed and 70 ul of 5k mPEG-silane (Sigma #JKA3037-1G, dissolved in 95% EtOH to a final concentration of 25 mg/ml) was added to each well. The plates were sealed and incubated overnight at room temperature. The mPEG-silane was removed, the wells were washed once with 95 % EtOH, then rinsed with copious amounts of MQ and dried completely in the fume hood.

The wells were then passivated by adding 40 ul of 20 mg/ml BSA (NEB) and incubated for at least one hour at room temperature. The BSA was removed and the wells were washed three times with Condensation Buffer (20 mM HEPES pH 7.5 at 20 °C, 0.2 mg/ml BSA, 10 % Glycerol, 5 mM DTT and 150 mM KCl). The chromatin stock was adjusted to a DNA concentration of 100 ng/ul (unless otherwise stated) and 150 mM KCl. The PRC1 complex was diluted in Condensation buffer to a protein concentration equal to twice the final PRC1 assay concentration as stated. To induce condensation, 16 ul of the diluted PRC1 were combined with 16 ul of the salt-adjusted chromatin dilution in PCR tubes. The samples were incubated for 30 minutes at room temperature before being transferred to the 384-well plate and incubated for a further 60 minutes at room temperature, so that the first images were recorded 90 minutes after induction of condensation.

Images were recorded with a Nikon C1 scanning confocal microscope. GFP was excited with a 488 nm laser, Cy5 was excited with a 561 nm laser. Linear contrast adjustments were made with ImageJ. Where several micrographs are compared to each other, the same contrast settings were used for all micrographs.

For condensation assays in Fig. 2a, Fig. 4, Extended Data Fig. 1d and Extended Data Fig. 9, chromatin and PRC1 dilutions were prepared in Chromatin Buffer (10 mM TRIS-HCl pH 7.5 at 20 °C, 10 mM KCl, 1 mM DTT) and PRC1 buffer (20 mM HEPES-KOH pH 7.5 at 20 °C, 150 mM NaCl, 1 mM DTT), respectively. After adjusting BSA and salt concentration and combining PRC1 and chromatin, the final reaction contained the PRC1 and chromatin concentration stated in the figures and the following components: 20 mM HEPES-KOH, pH 7.5, 5 mM TRIS-HCl pH 7.5 at 20 °C, 90 mM KCl, 32.5 mM NaCl, 0.2 mg/ml BSA (NEB), 1mM DTT.

DIC images and fluorescence wide field images were recorded with a Leica DMI8 imaging system equipped with a 4.2 MP (2k x 2k) sCMOS monochromatic K8 camera and a 63x oil immersion objective with numerical aperture of 1.3. DIC images in Extended Data Fig. 9 were recorded with a Leica DMI8 imaging system equipped with a 4.2 MP (2k x 2k) sCMOS monochromatic DFC 9000GT camera and a 40x dry objective with a numerical aperture of 0.8.

Condensate quantification was done using ImageJ. A threshold was set for each image to segment condensates in the frame using the Cy5 signal. Condensates overlapping the frame edges were excluded. Then the total area of all condensates in the frame was calculated.

### Chromatin ubiquitylation assay

The salt concentration of the chromatin stock was adjusted to 100 mM KCl. The nucleosome equivalent concentration of chromatin arrays was calculated by measuring the molar DNA concentration and assuming that one DNA molecule is populated by 20 nucleosomes. The reaction mixture included 750 nM (Fig. 1d) or 1000 nM (Extended Data Fig.1d) of chromatin (nucleosome equivalent concentration), 500 nM PRC1^C8^ or RING1b-BMI1 dimer, 100 nM hUBA1, 500 nM UBCH5C and 50 μM ubiquitin in Ub-Buffer (25 mM HEPES-KOH pH 7.5 at 20 °C, 100 mM KCl, 3 mM MgCl and 2 mM DTT) and started by adding ATP to a final concentration of 3 mM. 15 ul reactions were incubated at 30 °C for 45 minutes or the indicated time points (Extended Data Fig 1d). The reaction was stopped by adding 5 ul of 4X NuPage LDS-loading dye (Thermo Scientific cat #NP0008) supplemented with 5 % 2-mercaptoethanol. The samples were separated on a 4-12% NuPage gel (Thermo Scientific cat #NP0321BOX) using MES buffer (Thermo Scientific cat #NP0002) in the tank. The gels were then blotted onto a nitrocellulose membrane (Amersham, cat #GE10600002) in Tris-Glycine transfer buffer + 20 % Ethanol (v/v) for 90 minutes in the cold room at 310 mAmp in a blotting tank (BioRad). H2A was detected using anti-H2A primary antibodies (Merck Millipore Cat. # 07-146, 1:1000 titer) and HRP-conjugated secondary antibodies (Santa Cruz, cat #sc-2357, titer 1:5000).

### Sample preparation for cryo-electron tomography

The PRC1^C8^ complex (MBP-tagged CBX8) was combined with chromatinized ATOH1 at a final concentration of 1.6 μM PRC1^C8^ and 500 ng/ul DNA at a final salt concentration of 25 mM NaCl and 8.3 mM KCl. Samples were incubated at room temperature for 30 minutes. Just before freezing, 5 nm gold nanoparticles were added at a 1:6 ratio (v/v). 3.5 ul of sample was applied to a Quantifoil grid (R1.2/1.3 on 200 copper mesh, Quantifoil cat #N1-C14nCu20-01) and vitrified in liquid ethane using the Vitrobot plunge freezer (Thermo Scientific) with the following settings: Temperature 4°C, blot force -3, blot time 4 seconds, humidity 100%. The final sample composition after addition of gold was 1330 nM PRC1^C8^ and 3500 nM chromatin (estimated nucleosome concentration) in 3.5 mM HEPES-KOH pH 7.5, 6.8 mM TRIS-HCl PH 7.5, 21 mM NaCl, 7 mM KCl, 0.8 mM DTT. The sample of chromatin in absence of PRC1 was prepared the same but instead of adding PRC1 an equivalent volume of the PRC1 buffer was added.

### Low magnification cryo-EM image collection

The images in Extended Data Fig. 3a were collected just before cryo-ET data collection using the Titan Krios (Thermo Fisher) at an acceleration voltage of 300 keV and are of the same grid from which tomograms were collected. The images in Extended Data Fig. 4a were recorded using a Talos Arctica TEM (Thermo Fisher) at an acceleration Voltage of 200 keV.

### Cryo-electron tomography data collection and processing

The data was collected with a Titan Krios electron microscope (Thermo Fisher) at 300 keV acceleration voltage using a Gatan K2 (+PRC1 sample) or K3 (-PRC1 sample) Summit camera (Gatan). A tilt series was acquired ranging from -60 to 60 degrees with 3 degree increments and a nominal defocus of -2.5 μm. A dose symmetric collection scheme was followed as described previously^57^. Chromatin in the presence of PRC1 was images at pixel size of 1.32 Angstrom. The chromatin samples in the absence of PRC1 were collected in super resolution mode with a nominal pixel size of 1.632 Angstrom (0.815 Angstrom super resolution pixel size). The total dose per tomogram was 144.32 e/A^2^ (+PRC1) and 160.72 e/A^2^ (-PRC1). Four frames were collected per tilt (dose per frame 0.88 e/A^2^ for the +PRC1 sample and 0.98 e/A^2^ for the -PRC1 sample). The movies were motion corrected using motioncor2^58^. The motion corrected images were combined into stacks and further processed with Imod^59^ version 4.9.9. Imaging artefacts were identified and removed with Imod’s Ccderaser function. Tilts were aligned using the gold fiducial markers and the final aligned stack was binned by a factor of 4. The defocus was estimated using emClarity^60^ and the estimated defocus was used in Imod for CTF correction. The final tomogram was calculated using Imod’s implementation of weighted back projection. For visualisation, the tomogram was denoised using Topaz^50^. The denoised tomogram was only used for visualisation (Figure 1 and Movie S1). The original non-denoised tomograms were used for all further processing, including template matching and subtomogram averaging.

### Template matching, subtomogram averaging and modelling the chromatin structure of condensates from cryo-electron tomograms

Two tomograms were selected for further processing (Tomogram #1 and Tomogram #2 in the following). To avoid user bias, we used a template matching algorithm^61^ with the structure of a single nucleosome as template (EMD-8140^61^) to identify initial positions and orientations for each nucleosome. The pixel size of the template was adjusted using emClarity (version 1.0.0) to match the unbinned pixel size of the tomogram^60^. The template was then subjected to a low-pass filter of 30 Angstrom and binned by a factor of four to match the binned tomogram. Template matching was done with the Matlab implementation of Dynamo (version v-1.1.514)^61^ using the “dynamo_match” function. The template was masked with a tight-fitting mask with smooth edges. The scanning range was set to 360 degrees with 10 degree steps. In-plane rotation was also scanned over 360 degrees with 10 degrees steps. Particles that passed a cross-correlation cut-off of 0.17 (Tomogram #1), 0.19 (Tomogram #2), 0.24 (Tomogram #27), 0.23 (Tomogram #41) and 0.25 (Tomogram #49) were selected for further analysis. Obvious false positives were removed manually.

The particles were cropped from the tomogram with a box size of 36 pixels. Nucleosome position and orientation was refined over three rounds of subtomogram averaging^61^. As an initial template for subtomogram averaging, we used an average from all cropped particles after template matching, masked with a tight-fitting mask with smooth edges generated in the Dynamo mask editor. Specific settings for the different rounds of subtomogram averaging are detailed in Table 2. The resulting average structure of a nucleosome from the subtomogram averaging was then used to populate a volume the size of the tomogram with nucleosomes at the determined positions and orientations. The graphic depiction of the final model (Figure 1G) was generated using the Dynamo Matlab implementation^61^.

**Table 2:**
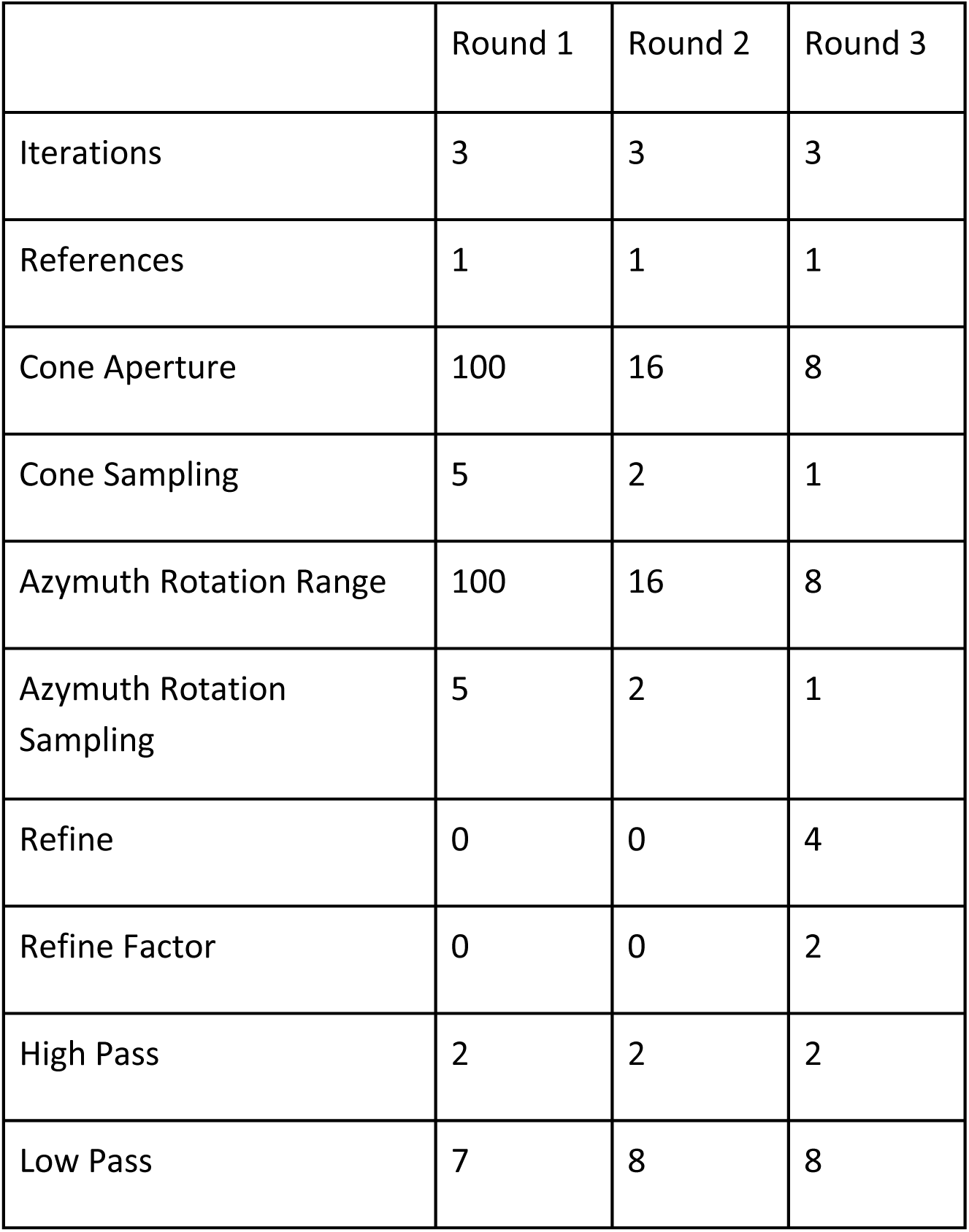
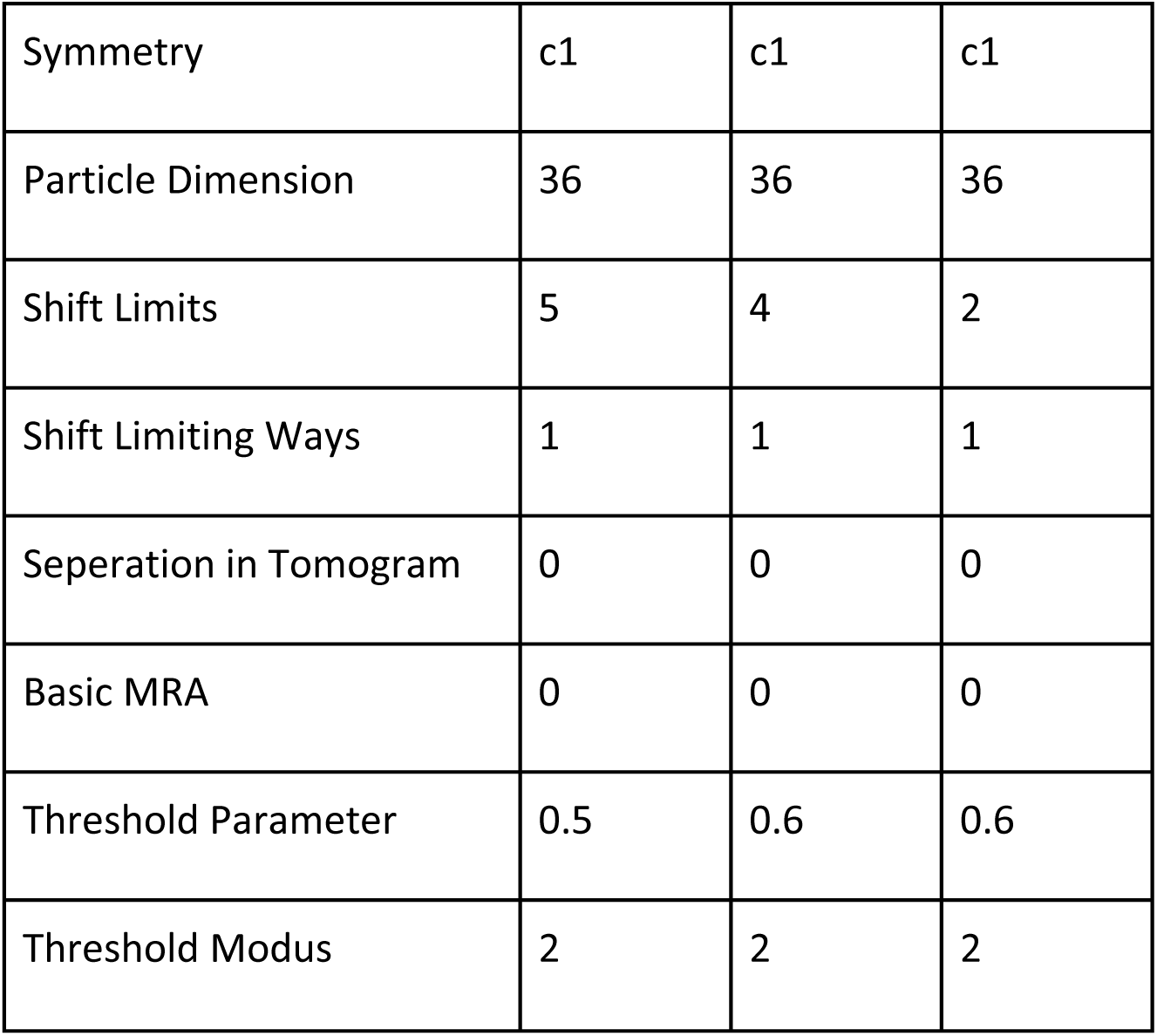
Settings for subtomogram averaging in Dynamo.

|  | Round 1 | Round 2 | Round 3 |
| --- | --- | --- | --- |
| Iterations | 3 | 3 | 3 |
| References | 1 | 1 | 1 |
| Cone Aperture | 100 | 16 | 8 |
| Cone Sampling | 5 | 2 | 1 |
| Azimuth Rotation Range | 100 | 16 | 8 |
| Azimuth Rotation Sampling | 5 | 2 | 1 |
| Refine | 0 | 0 | 4 |
| Refine Factor | 0 | 0 | 2 |
| High Pass | 2 | 2 | 2 |
| Low Pass | 7 | 8 | 8 |
| Symmetry | c1 | c1 | c1 |
| Particle Dimension | 36 | 36 | 36 |
| Shift Limits | 5 | 4 | 2 |
| Shift Limiting Ways | 1 | 1 | 1 |
| Seperation in Tomogram | 0 | 0 | 0 |
| Basic MRA | 0 | 0 | 0 |
| Threshold Parameter | 0.5 | 0.6 | 0.6 |
| Threshold Modus | 2 | 2 | 2 |

### Analysis of exclusion volume

Exclusion volumes for spherical molecules of various radii were calculated using 3V^26^ as follows. First, the positional and rotational coordinates of the nucleosomes within Tomogram #1 were determined by template matching followed by subtomogram averaging, as described above. Next, the positional and rotational coordinates of the nucleosomes were tabulated from within a section of 300 pixels x 300 pixels x 150 pixels (corresponding to 158.4 nm x 158.4 nm x 87.0 nm at the x, y and z axes, respectively) centred on the pixel at location (650,650,80), which was large enough to include hundreds of nucleosomes but yet sufficiently small to carry out the subsequent computational analysis. The dimensions of the resulting table were converted to Angstrom by multiplying with a tomogram pixel size of 5.28 Angstrom/pixel. Next, the table including nucleosome positions was used by the Dynamo function ‘dtchimera’ to generate a chimera cmd script that placed a pdb structure of a single nucleosome (PDB 1KX4^63^) at the position and orientation defined for each nucleosome in the table. At the end of this process, each of the nucleosomes within the tomogram section is represented by the pdb coordinates of the nucleosome template. The resulting model was saved as a pdb file and used as input for 3V^26^. Varying probe radii ranging from 2 to 20 nm were used.

### Hydrodynamic radius calculation using protein structures

The hydrodynamic radii of various proteins (Figure 1I) were calculated using the HullRad^39^ web server (http://52.14.70.9/). The PDB codes for the structures used are: 6LTJ (BAF^45^), 1MUH (Tn5^40^), 7O4J (PolII-PIC^47^), 6C24 (PRC2.2^64^), 8GXS (PolII-Mediator^65^). 6LTJ, 6C24, 8GXS and 7O4J are only partial models as not all residues were assigned, therefore the calculated hydrodynamic radius represents an estimate of the minimal complex size, while the actual size of these complexes could be larger.

### Analysis of nucleosome-nucleosome orientation and distances

Nucleosome-nucleosome orientation was classified into face-face, face-side and side-side, as described in^22^. The orientations were calculated from the output table of the template matching and subtomogram averaging process in Dynamo^61^ using the Matlab script ‘calculate_orientation.m’. The nucleosome-nucleosome orientation plot (Fig. S1A) was generated using the ‘plot_edges’ function from the python notebook NCP_orientation_analysis.ipyn. Both scripts are available on Github (https://github.com/MichaUckelmann/Chromatin-Structure-Analysis). Centre-centre distances between neighbouring nucleosomes were calculated using the Matlab function ‘knnsearch’. Double-counted distances were removed before further analysis.

### Cryo-light microscopy

The samples were prepared as described for cryo-electron tomography, without gold nanoparticles and in a buffer containing 100 mM KCl. The vitrified grids were imaged using a ZEISS LSM900 Airyscan2 with a Linkam CMS196V Cryo stage.

### Fluorescence recovery after photobleaching (FRAP)

For FRAP of PRC1^C8^ that was fluorescently labelled using a GFP-CBX8, samples and plates were prepared and images were recorded as described above for chromatin condensation assays. 1 μM PRC1^C8^ with an N-terminal GFP tag was used. The FRAP experiments were set up with the NIS-Elements software (Nikon). Regions of interest (ROIs) were defined and a single image was recorded before bleaching. Then the ROI was bleached using 488 nm (GFP) and 561 (Cy5) lasers. The bleaching time and laser power was set so that approximately 80 % of fluorescence signal within the ROI was quenched. Recovery was measured over 424 seconds, recording a total of 13 images. The data was analysed with ImageJ, the mean pixel intensity of the bleached ROI was quantified for each timepoint.

For FRAP of PRC1^C8^ that was fluorescently labelled using spared labelling of lysine residues using Atto-488, samples and plates were prepared and images were recorded as described above for chromatin condensation assays. To bring both the untagged PRC1^C8^ and the MBP-tagged PRC1^C8^ (i.e. an N-terminal MBP tag on CBX8) to the same concentration, the stock solutions of the Atto-488-labelled untagged PRC1^C8^ was adjusted to 8 μM PRC1^C8^, 150 mM NaCl and 25mM HEPES pH7.5. Both the PRC1^C8^ stocks, with or without the MBP tag, were then diluted to 4 μM protein in a condensation buffer (25 mM HEPES pH7.5, 150 mM KCl, 0.2 mg/mL BSA, 2 mM DTT). 8 μl of 4 μM PRC1 solution and 8 μl of 100 ng/μl chromatin (DNA concentration) were combined in a well to induce condensation, at final concentrations of 2 μM PRC1^C8^, with or without the MBP tag, 50 ng/μl chromatin (DNA concentration) and with a final buffer composition of 25 mM HEPES pH7.5, 37.5 mM NaCl, 112.5 mM KCl, 0.1 mg/mL BSA and 1 mM DTT. Regions of interest (ROI) were defined and a single image was recorded before bleaching. Then the ROI was bleached using 488 nm (Atto-488) and 561 nm (Cy5) lasers. The bleaching time and laser power were set so that approximately 80 % of the fluorescence signal within the ROI was quenched. Recovery was measured over 24 minutes, with a frequency of ∼ 1 image/minute. The data was analysed with NIS-Elements software.

### Single molecule confocal microscopy

Cy5 labelled reconstituted chromatin (200 nM nucleosome concentration) and GFP labelled PRC1^C8^ at the concentration indicated in the figure were combined in assay buffer (25 mM HEPES-KOH pH 7.5, 100 mM KCl, 15 mM NaCl, 1 mM DTT, 0.1 mg/ml BSA). Samples were immediately loaded into a custom-made silicone well plate with a 70 x 80 mm glass coverslip (ProSciTech, Kirwan, QLD, Australia). Plates were analysed at room temperature on a custom setup based on a Zeiss Axio Observer microscope (Zeiss, Oberkochen, Germany). Illumination is provided by a 488 nm and a 561 nm laser beams, cofocussed in the sample volume using a ×40 magnification, 1.2 Numerical Aperture water immersion objective (Zeiss, Oberkochen, Germany). This creates a very small observation volume in solution (∼1 fL), through which fluorescent proteins diffuse and emitting light in specific wavelengths as their fluorescent tags are excited by the laser beams. Light emitted by the fluorophores is split into GFP and mCherry channels by a 560 nm dichroic mirror. The fluorescence of GFP is measured through a 525/50 nm band-pass filter and the fluorescence of mCherry is measured through a 580 nm long-pass filter. Fluorescence is detected by two photon counting detectors (Micro Photon Devices, Bolzano, Italy). Photons of the two channels are recorded simultaneously in 1 ms time bins by a custom Lab-VIEW 2018 program (National Instruments)^66^. The data were analysed using a custom python script (Spyder version 4.1.5) that automatically detects peaks, as described in^67,68^.

### DNA binding assays

DNA binding was assayed using a 24 bp DNA (sequence see Table 1) with Fluorescein attached to the top strand. The probe was protected from light wherever possible. The probes were synthesised and delivered as duplexes (IDT). Probes were dissolved at a concentration of 5 mM DNA in milliQ water. The probes were then diluted to 4 μM in annealing buffer (20 mM HEPES-KOH, 150 mM NaCl) and heated to 95 °C for 5 minutes. The probe was then left at room temperature for at least 1 hour to anneal before being diluted to 20 nM in 20 mM HEPES-KOH pH 7.5 at 20 °C, 150 mM NaCl, 1 mg/ml BSA (NEB cat #B9000S), 0.1% Tween20, 1 mM DTT. Serial protein dilutions of the PRC1^C8^ and RING1b-BMI1 complexes were prepared in Protein Dilution Buffer (20 mM HEPES-KOH, 150 mM NaCl, 1 mM DTT) ranging from 8000 nM to 3.9 nM (Fig. 3) and from 7200 nM to 7 nM (Fig. 4). 20 ul of probe were mixed with 20 ul of the respective protein dilution and transferred to a 384-well plate. The samples were incubated for 30 minutes in the dark at room temperature and then read using a Pherastar plate reader (BMG Labtech). The fluorescence anisotropy signal was normalised and the curves were fitted with a specific binding model with Hill slope (GraphPad Prism).

### Crosslinking mass spectrometry (XL-MS)

Sample preparation, processing and data analysis was identical for samples with and without chromatin. Reconstituted chromatin (if used) was dialysed overnight against 1 litre of XL buffer (25 mM HEPES-KOH pH 7.5 at 20 °C, 150 mM NaCl, 1 mM DTT). The chromatin was combined with PRC1^C8^ in a 1:1.5 molar ratio (Nucleosomes:PRC1^C8^). Specifically, the nucleosome equivalent concentration of the chromatin array was calculated from the measured DNA concentration assuming 18 nucleosomes per DNA molecule. 1.5 μM PRC1 and 1 μM nucleosome equivalent concentration of chromatin arrays were then combined in XL buffer and incubated for 30 minutes at room temperature. Crosslinking was done as described before^69^. The bis(sulfosuccinimidyl)suberate (BS3) crosslinker was added to a final concentration of 500 μM and crosslinking proceeded for 20 minutes at room temperature at a reaction volume of 15 μl. The reaction was stopped by the addition of Tris-HCl pH 8 at 20 °C to a final concentration of 125 mM. The samples were then diluted to a volume of 100 µL using a buffer containing 50 mM Tris pH 8 at 20 °C and 150 mM NaCl. TCEP was added to a final concentration of 10 mM and the samples were incubated at 60 °C for 30 min. Chloroacetamide was added to a final concentration of 40 mM and the samples were incubated in the dark for 20 min. The samples were then digested using trypsin (Promega cat #V5280) at 37 °C overnight. The digest was stopped by adding formic acid to a final concentration of 1% v/v. The digested samples were purified using 100 µl ZipTip pipette tips (Merck cat #ZTC18S960) according to the manufacturer’s instructions. Samples were then concentrated to ∼5 μL using a SpeedVac vacuum centrifuge and diluted with 20 µL Buffer A (0.1% v/v formic acid).

The peptides were analyzed by online nano-high-pressure liquid chromatography (UHPLC) electrospray ionization-tandem mass spectrometry (MS/MS) on an Q Exactive Plus Instrument connected to an Ultimate 3000 UHPLC (Thermo-Fisher Scientific). Peptides reconstituted in 0.1% formic acid were loaded onto a trap column (Acclaim C18 PepMap 100 nano Trap, 2 cm × 100 μm I.D., 5-μm particle size and 300-Å pore size; Thermo-Fisher Scientific) at 15 μL/min for 3 min before switching the precolumn in line with the analytical column (Acclaim C18 PepMap RSLC nanocolumn, 75 μm ID × 50 cm, 3-μm particle size, 100-Å pore size; Thermo-Fisher Scientific). The separation of peptides was performed at 250 nL/min using a non-linear acetonitrile (ACN) gradient of buffer A (0.1% formic acid) and buffer B (0.1% formic acid, 80% ACN), starting at 2.5% buffer B to 42.5% over 95 min. Data were collected in positive mode using a Data Dependent Acquisition m/z of 375–2000 as the scan range, and higher-energy collisional dissociation (HCD) for MS/MS of the 12 most intense ions with z 2–5. Other instrument parameters were: MS1 scan at 70,000 resolution, MS maximum injection time 118 ms, AGC target 3E6, ion intensity threshold of 4.2e4 and dynamic exclusion set to 15 s. MS/MS resolution of 35000 at Orbitrap with the maximum injection time of 118 ms, AGC of 5e5 and HCD with collision energy = 27%.

For the data analysis, Thermo raw files were analysed using the pLink 2.3.4 search engine^70^ to identify crosslinked peptides, searching against the sequences of RING1b, BMI1, CBX8, H2A, H2B, H3 and H4. The default settings for searches were used. N-terminal acetylation and methionine oxidation were used as variable modifications and carbamidomethyl on cysteines as a fixed modification. False discovery rates of 1% for peptide spectrum match level were applied by searching a reverse database. Crosslinked peptides were further analysed using the crissrosslinkeR package^71^. Specifically, peptides were retained by crissrosslinkeR only if they passed a p-value cutoff of 0.05 or were present in at least two of three replicates. Subsequent visualisation of retained peptide was carried out with xiNET^72^.

### Generation of Cbx8 KO mESC lines using CRISPR/Cas9

Paired sgRNAs were designed to delete exons 1–4 of the murine *Cbx8* gene. The Golden Gate Cloning method was used to clone the sgRNAs (sequence below) into the lentiguide-mCherry-Cas9 plasmid^73,74^. 2 million mESCs were transfected with 1 μg of each plasmid carrying sgRNAs-mCherry-Cas9, using electroporation (Neon™ Transfection System MPK5000). The following day, mCherry-positive mESCs were sorted by FACS and plated on a 10 cm dish at a very low density for single-cell clone picking. After 5-6 days, individual colonies (derived from single cells) were picked, expanded, and genotyped using genomic PCR to identify homozygous/biallelic deletions of Cbx8 KO mESC colonies. Selected Cbx8 KO mESC lines were further confirmed by western blot for CBX8 (Cell Signalling, CBX8 (D2O8C), cat # 14696S, titer 1:50 (Fig. 4a)) and HRP-conjugated secondary antibodies (Santa Cruz, cat #sc-2357, titer 1:10000).

### Cbx8 sgRNA sequences (5’ and 3’ sgRNAs)

CBX8-5’Fw: CACCTGCGAATGCGCCGCTTCAGG

CBX8-5’Rv: AAACCCTGAAGCGGCGCATTCGCA

CBX8-3’Fw: CACCCTCTATGGCCCCAAAAAGCG

CBX8-3’Rv: AAACCGCTTTTTGGGGCCATAGAG

### Cbx8 genotyping primers

Deletion_Fw: GCCTTCTGGTGCAGCTAAGT

Deletion_Rv: GACGTCAGCGGGAGAGTATT

Internal_Fw: CACCAAATGAATGCTCCAAA

Internal_Rv (same as the Deletion_Rv): GACGTCAGCGGGAGAGTATT

### Mouse embryonic stem cell culture

Wildtype and *Cbx8* knockout mES cells were grown on gelatinized culture dishes in DMEM, 20 % FBS, 1x non-essential amino acids (Gibco #11140-050), 1x Glutamax (Gibco #35050-061), PenStrep 100 u/ml (Thermo Fischer), 0.5 x EmbryoMax 2-Mercaptoethanol (Merck Millipore #ES-007-E), 2.5 μg/ml Plasmocin (Invivogen), 1000 U/ml ESGRO Leukemia Inhibitory Factor (LIF) (Merck Millipore cat #ESG1107). For ATAC-seq experiments with the reporter-integrated mECS line, the cells were treated for 6 days with 1 µg/ml Doxycycline (passaged every 48 h) before ATAC-seq was performed as described below.

### Mouse embryonic stem cell differentiation ahead of ChIP-seq and ATAC-seq

Differentiation was induced by seeding cells at a density of 0.3×10^6^ cells per well in 6-well plates (for ATAC-seq) or at 1.5×10^6^ cells per 10-cm culture dish (for ChIP-seq) in media containing 1 μM all-trans retinoic acid (RA,Sigma-Aldrich R2625-50MG) and no LIF. Cells were differentiated for 48 hours and the media was changed after 24 hours. Control cells were treated with a DMSO volume equivalent to the RA volume in the differentiating cells. After 48 hours, the cells were harvested, washed once with PBS, counted and used immediately in either ChIP-seq or ATAC-seq experiments.

### Mouse embryonic stem cell culture for ChIP-qPCR

TetR-CBX8 reporter-integrated mESCs were cultivated without feeders in high-glucose-DMEM (Corning 10-013-CV) supplemented with 13.5% fetal bovine serum (Corning 35-015-CV), 10 mM HEPES pH 7.4 (Corning, 25-060-CI), 2 mM GlutaMAX (Gibco, 35050-061), 1 mM Sodium Pyruvate (Corning 25-000-Cl), 1% Penicillin/Streptomycin (Sigma, P0781), 1X non-essential amino acids (Gibco, 11140-050), 50 mM β-mercaptoethanol (Gibco, 21985-023) and recombinant LIF. Cells were incubated at 37 °C and 5 % CO2 and were passaged every 48 h by trypsinization in 0.25 % 1x Trypsin-EDTA (Gibco, 25200-056). To reverse TetR-CBX8 binding, 1 µg/ml Doxycycline (Sigma, D9891) was added to cell culture medium for 6 hours.

### ChIP qPCR

25 x 10^6^ reporter-integrated mESCs were collected, washed once in 1x PBS and cross-linked for 7 min in 1 % formaldehyde. The crosslinking was quenched by addition of 125 mM glycine and incubated on ice. The cross-linked ESCs were pelleted by centrifugation for 5 min at 1200 g at 4 °C. Nuclei were prepared by washes with NP-Rinse buffer 1 (10 mM Tris pH 8.0, 10 mM EDTA pH 8.0, 0.5 mM EGTA, 0.25 % Triton X-100) followed by NP-Rinse buffer 2 (10 mM Tris pH 8.0, 1 mM EDTA, 0.5 mM EGTA, 200 mM NaCl). Afterwards, the nuclei were washed twice with shearing buffer (1 mM EDTA pH 8.0, 10 mM Tris-HCl pH 8.0, 0.1 % SDS) and subsequently resuspended in 900 µL shearing buffer including 1x protease inhibitors Complete Mini cocktail (Roche). Chromatin was sheared by sonication in 15 ml Bioruptor tubes (Diagenode, C01020031) with 437.5 mg sonication beads (Diagenode, C03070001) for 6 cycles (1 min on and 1 min off) on a Bioruptor Pico sonicator (Diagenode). ChIP lysates equivalent to 50 ug DNA were incubated in 1x IP buffer (50 mM HEPES/KOH pH 7.5, 300 mM NaCl, 1 mM EDTA, 1% Triton X-100, 0.1% DOC, 0.1% SDS), and 1.5 ul of FLAG M2 antibody (Sigma Aldrich Sigma F1804) overnight. Antibody-bound chromatin was captured using Dynabeads protein G beads (Thermofisher #10004D) for 4 h at 4 °C. ChIPs were washed 5x with 1x IP buffer (50 mM HEPES/KOH pH 7.5, 300 mM NaCl, 1 mM EDTA, 1% Triton-X100, 0.1 % DOC, 0.1 % SDS), followed by 3x washes with DOC buffer (10 mM Tris pH 8, 0.25 mM LiCl, 1 mM EDTA, 0.5 % NP40, 0.5 % DOC) and 1x with TE/50 mM NaCl. ChIP DNA was eluted twice with elution buffer (1 % SDS, 0.1 M NaHCO_3_) at 65 °C for 20 min, and subsequently treated with RNase A (60 ug final, Invitrogen) for 30 min at 37 °C, and Proteinase K (15ug, NEB) for 3 h at 55 °C and crosslinks were reversed overnight at 65 °C. The following day, ChIP samples and corresponding inputs were purified by AMPure XP beads (Beckman Coulter A63880).

ChIP-qPCR primers:

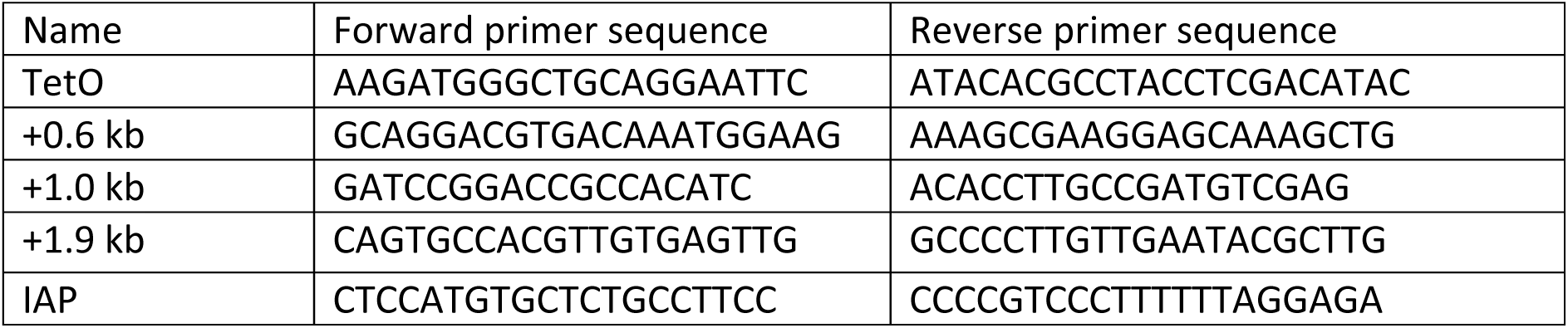

### ChIP-seq

ChIP-seq was done as described previously^75^. 1.5 μg of CBX8 antibody (Cell Signalling, CBX8 (D2O8C), cat # 14696S) and 3 μg of H3K27me3 antibody (Tri-Methyl-Histone H3 (Lys27) (C36B11), cat #35861SF) were used. Libraries were prepared using NEBNext ultra II DNA library kit for Illumina (NEB Biolabs) according to the manufacturer instructions. The resulting libraries were assessed for quality on a High Sensitivity D1000 Screen Tape (Agilent) and were sequenced using an Illumina Novaseq 6000 sequencer (Genewiz/Azenta).

### Data processing for Chip-seq

The reads were quality-trimmed and adapters were removed using Trim Galore!, a wrapper script to cutadapt^76^, in paired end mode and using default settings. The reads were then aligned to the mouse mm10 genome build using bowtie2 (version 2.3.5)^77^ with the option “very-sensitive”. The data was reduced to only properly paired reads using “samtools view” (Samtools version 1.9) with the flag “-f 3”. PCR duplicates were removed using the RemoveDuplicates function from Picard Tools (version 2.19.0). Read mates were fixed using samtools fixmate. BigWig files were calculated using BamCoverage (Deeptools version 3.5.2) with CPM normalisation.

Peaks were called with Macs2 (version 2.1.1)^78^ callpeak function, using the input sample as control. For the CBX8 ChIP-seq data set, default settings with a q-value cut-off of 0.05 were used. For H3K27me3 ChIP-seq data set, broad mode was used with a q-value and broad cutoff of 0.001. Peaks overlapping with ENCODE blacklisted regions^79^ were removed.

### ATAC-seq

ATAC-seq was done using a commercial kit (Diagenode Cat.# C01080002) according to the instructions of the manufacturer. The resulting libraries were assessed for quality control on a High Sensitivity D1000 Screen Tape (Agilent) and libraries were sequenced using an Illumina Novaseq 6000 sequencer (Genewiz/Azenta).

### Data processing ATAC-seq

ATAC-seq data was processed as described previously^80^. Specifically, reads were quality-trimmed and adapters were removed using Trim Galore!, a wrapper script to cutadapt^76^, in paired end mode using default settings. The reads were then aligned to the mouse mm10 genome build using bowtie2 (version 2.3.5)^77^ with the option “very-sensitive”. Reads were sorted and indexed using Samtools (version 1.9). Mitochondrial reads were removed using a python script from Harvard Bioinformatics (available at https://github.com/harvardinformatics/ATAC-seq). The data was reduced to only properly paired reads using “samtools view” with the flag “-f 3”. The library complexity was estimated and the data sets were subsampled to reach a similar complexity as described previously^80^. Data shown in Fig. 5h and Extended Data Fig 8b were not subsampled because complexity was nearly identical. PCR duplicates were removed using the RemoveDuplicates function from Picard Tools (version 2.19.0). Read mates were fixed using samtools fixmate. BigWig files were calculated using BamCoverage (Deeptools version 3.5.2) with CPM normalisation For peak calling the bam files were converted to BEDPE files and the Tn5 shift was corrected by running a bash script provided at https://github.com/reskejak/ATAC-seq (bedpeTn5Shift.sh). Files were then converted to minimal bed format and peaks were called using Macs2 (version 2.1.1)^78^ callpeak function in broad mode with broad-cutoff set to 0.05. Peaks overlapping with ENCODE blacklisted regions^79^ were removed. Consensus peaks for each condition were defined as the intersect of peaks from both biological replicates. Venn diagrams were generated using the ChIPPeakAnno package^81^.

